# Characterisation of the Ubiquitin-ESCRT pathway in Asgard archaea sheds new light on origins of membrane trafficking in eukaryotes

**DOI:** 10.1101/2021.08.17.456605

**Authors:** Tomoyuki Hatano, Saravanan Palani, Dimitra Papatziamou, Diorge P. Souza, Ralf Salzer, Daniel Tamarit, Mehul Makwana, Antonia Potter, Alexandra Haig, Wenjue Xu, David Townsend, David Rochester, Dom Bellini, Hamdi M. A. Hussain, Thijs Ettema, Jan Löwe, Buzz Baum, Nicholas P. Robinson, Mohan Balasubramanian

**Author notes:** Correspondence to Mohan Balasubramanian-; Nick Robinson-; Buzz Baum. These authors contributed equally to this work and are listed alphabetically (Saravanan Palani, Dimitra Papatziamou, Diorge P. Souza, Ralf Salzer, Daniel Tamarit).

## Abstract

The ESCRT machinery performs a critical role in membrane remodelling events in all eukaryotic cells, including in membrane trafficking, membrane repair, cytokinetic abscission, in viral egress, and in the generation of extracellular vesicles. While the machinery is complex in modern day eukaryotes, where it comprises dozens of proteins, the system has simpler and more ancient origins. Indeed, homologues of ESCRT-III and the Vps4 ATPase, the proteins that execute the final membrane scission reaction, play analogous roles in cytokinesis and potentially in extracellular vesicle formation in TACK archaea where ESCRT-I and II homologues seem to be absent. Here, we explore the phylogeny, structure, and biochemistry of homologues of the ESCRT machinery and the associated ubiquitylation system found in genome assemblies of the recently discovered Asgard archaea. In these closest living prokaryotic relatives of eukaryotes, we provide evidence for the ESCRT-I and II sub-complexes being involved in the ubiquitin-directed recruitment of ESCRT-III,_as it is in eukaryotes. This analysis suggests a pre-eukaryotic origin for the Ub-coupled ESCRT system and a likely path of ESCRT evolution via a series of gene duplication and diversification events.

## INTRODUCTION

The ESCRT (Endosomal Sorting Complex Required for Transport) machinery is composed of several protein complexes and associated accessory proteins, including the ESCRT-0, −I, −II, −III subcomplexes, Vps4, and ALIX/Bro1^1–18^ (Figure S1). These proteins act in sequence to bind, deform and cut membranes during membrane trafficking^2–5^, cell division^19,20^, viral egress^21,22^ and in other important topologically similar membrane remodelling events in eukaryotes^23–39^. When driving the formation of multivesicular bodies (MVBs), where the role of ESCRT in trafficking has been best characterised, the ESCRT machinery is recruited to endosomal membranes by ubiquitylated transmembrane proteins that are targeted to vesicles^40^. The ubiquitin (Ub) moiety is recognised by ESCRT-0 and −I subcomplexes, which bind the ubiquitin β-grasp fold^9,14,41,42^. ESCRT-I proteins together with the ESCRT-II sub-complex then corral these ubiquitylated transmembrane proteins into membrane domains^9,11^. Finally, the ESCRT-II sub-complex nucleates the local formation of ESCRT-III co-polymers which, through action of the Vps4 ATPase, undergo structural changes to drive membrane invagination and scission to release vesicles containing the Ub-marked transmembrane proteins^43–53^.

ESCRT-I, −II, −III components are found conserved across the eukaryotic lineages, pointing to this machinery being present in the last eukaryotic common ancestor (LECA; Note ESCRT-0 is only encountered in Opisthokonta)^54^. Homologues of ESCRT-III and Vps4 are coded by the genomes of many archaeal species^55–60^, and shown to function in archaeal membrane remodelling during cytokinesis and virus release. More recently, PspA and Vipp1 have been recognized as bacterial ESCRT-III related proteins^61–63^. These observations suggest that a subset of ESCRT components have more ancient evolutionary origins. Furthermore, the identification of a full complement of ubiquitin and its activating enzymes (E1, E2, and E3) in some archaeal species has provided evidence for ubiquitylation cascades functioning in protein degradation in the archaeal ancestors of eukaryotes^58,64,65^.

This begs the question: when in evolution did ESCRT-I and ESCRT-II machineries arise, and when was ubiquitylation co-opted to regulate ESCRT? Metagenome assemblies of the recently discovered Asgard archaea, the closest living relatives of eukaryotes, have revealed that homologues of the entire ubiquitylation cascade, ESCRT-III (and Vps4), and components of the ESCRT-I and ESCRT-II subcomplexes are all encoded by the genomes of these archaea^57,58^. However, validating this conclusion in cells is currently very difficult, as only one Asgard member has been isolated and cultured^66^; and its growth rate, physiology and the lack of essential tools prevent its use as a cell biological model.

To circumvent these challenges, here we apply a diverse set of experimental approaches to characterise Asgard archaeal homologues of the eukaryotic Ub-ESCRT system, focusing on the ESCRT-I and ESCRT-II subcomplexes. Our analysis shows that, like its eukaryotic counterparts, the Asgard ESCRT-I subcomplex stably recognises ubiquitin. Furthermore, by carrying out a comprehensive two-hybrid analysis, we have been able to identify protein-protein interactions within and between the different ESCRT subcomplexes. Additionally, our data show that Asgard ESCRT subcomplexes have likely arisen through a process of gene duplication and diversification, prior to the evolution of more complex eukaryotic ESCRT assemblies. Taken together, this work reveals the presence of a genetically streamlined ubiquitin-associated multi-component ESCRT pathway that predates the emergence of the eukaryotic ESCRT machinery.

## RESULTS

### Phylogenetic analyses indicate that Asgard archaeal genomes encode homologues of most, but not all components of the Ub-ESCRT machinery

Many of the recently discovered Asgard archaeal genomes encode a wide array of so-called ‘eukaryotic signature proteins’ (ESPs) and appear unique amongst prokaryotes in possessing close homologues of most of the proteins that make up the ESCRT-I and −II complexes, together with ESCRT-III and Vps4 and homologues of ubiquitin and the associated ubiquitin-modification enzymes^57,58^. While these data suggest the possibility of some Asgard archaea possessing functional Ub-ESCRT membrane trafficking machinery, it is noticeable that Asgard genomes appear to lack a number of genes encoding proteins essential for ESCRT function in eukaryotes (Figure 1A). Thus, to put this hypothesis to the test, we began by generating a catalogue of proteins with homology to the components of the eukaryotic Ub-ESCRT pathway in distinct Asgard phyla shown in Figure 1A and Figure S2 and Figure S3.

**Figure 1.**
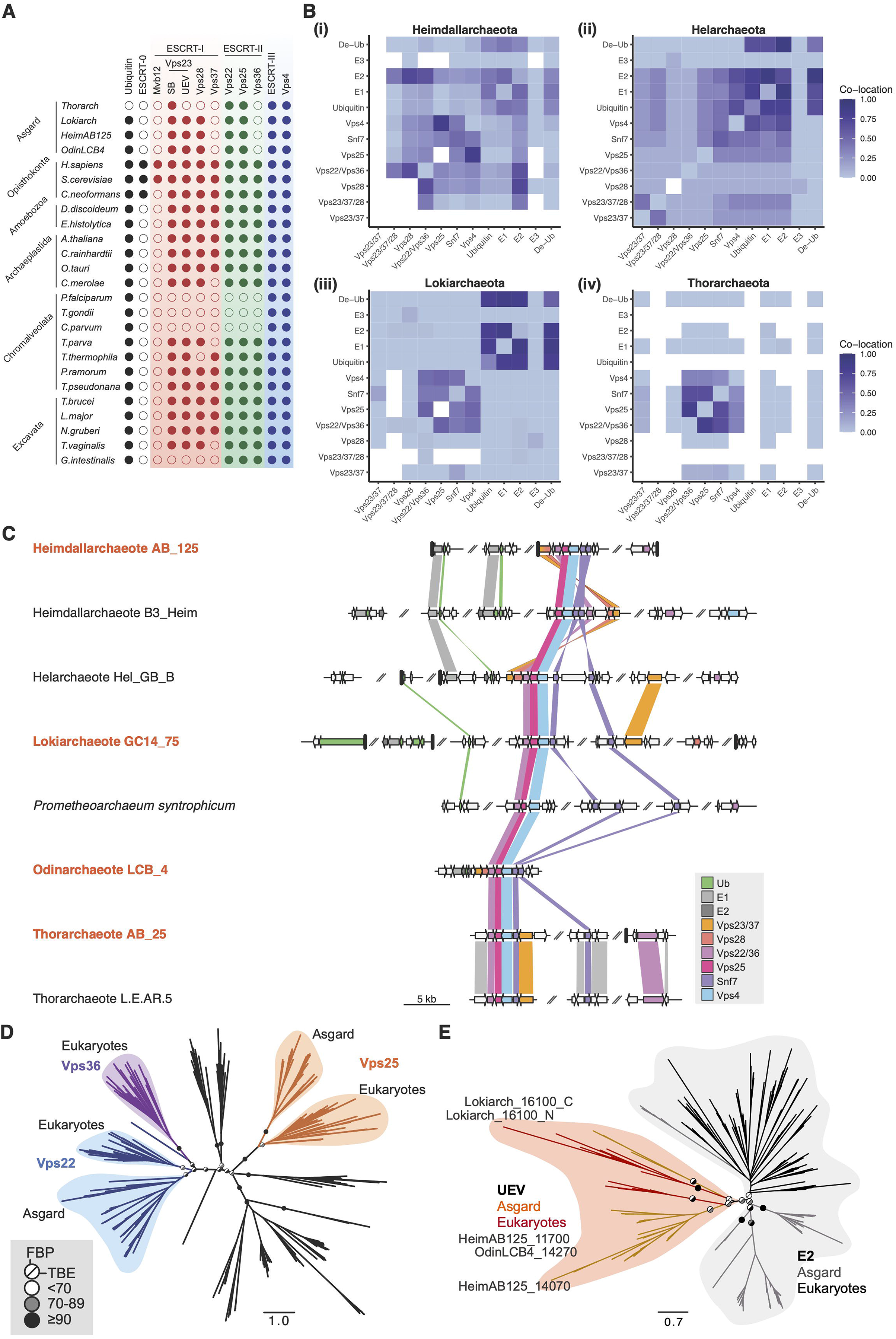
The ASGARD archaea, Lokiarchaeota, Heimdallarchaeota, Helarchaeota, and Odinachaeota, have genes that comprise the ubiquitin-ESCRT pathway. List of proteins in the Asgard archaea and eukaryotic Ubiquitin-ESCRT (Ub-ESCRT) pathway. Co-location of Ub/ESCRT protein-encoding genes in Heimdall-(i; 22 genomes), Hel-(ii; 9 genomes), Loki-(iii; 29 genomes) and Thorarchaeota (iv; 30 genomes). A colour gradient indicates the fraction of genomes in which a pair of genes was found to co-locate at less than 10 kb, and white cells indicate gene pairs found to never co-exist in any genome of the same phylum. Synteny plot of selected genomes. Arrows represent genes and are coloured if their products were annotated as containing diagnostic domains for Ub/ESCRT proteins (see Methods). Genes encoding Vps23/37 in the vicinity of the ESCRT gene cluster in Helarchaeote Hel_GB_B, Heimdallarchaeote B3_Heim and Odinarchaeote LCB_4 were only found to contain E2 domains, but were deemed Vps23/37 through alignment (Suppl. Fig. X). A gene encoding a fusion of Vps23/37 and Vps28 is colored as both orange and red. Genome regions are plotted at 2 kb of ubiquitin or ESCRT protein-encoding genes (coloured), or until a contig boundary (thicker vertical lines). Similarity lines indicate best-reciprocal BLAST-p (Altschul et al 1990 J Mol Biol 215: 403-410) hits with an e-value lower than 1e-5. The names of the organisms used for experimental analyses in later sections are marked in orange. Phylogenetic reconstruction of Vps22 and Vps36. Unrooted maximum likelihood phylogenetic tree of Vps22 (blue), Vps36 (purple) and Vps25 (orange) and outgroup (black) sequences. The tree was reconstructed using IQ-Tree under the LG+C60+R4+F model. Support values are only shown for the deeper branches connecting gene homologs and represent standard Felsenstein bootstrap proportions (upper left) or transfer-bootstrap expectation (TBE) (lower right) values based on 100 bootstrap pseudoreplicates. The full tree with all leaf and support labels is shown in Suppl. Figure X. Phylogenetic reconstruction of UEV domain-containing proteins and E2 ubiquitin-conjugating enzymes. Unrooted maximum likelihood phylogenetic tree of the UEV domain-containing proteins (orange, red) and E2 ubiquitin-conjugating proteins (gray, black) in Eukarya and Asgard archaea. The tree was reconstructed using IQ-Tree under the Q.pfam+C20+G4+F+PMSF model. Support values are only shown for the deeper branches, following the same pattern as in (D).

We focused on Asgard archaea that have metagenomic assemblies that are near complete based on them being assigned to small numbers of contigs and on estimates of genome coverage^57,58^. Within such genomes, as previously described^57,58^, we were able to identify genes coding for close homologues of ubiquitin, ubiquitin modifying enzymes, ESCRT-I components, ESCRT-II subunits, together with homologues of ESCRT-III and Vps4 (Figure 1A and Figure S3)^58,67^. These analyses strongly suggest that many Asgard archaea possess a *bona fide* eukaryote-like ESCRT-system.

Since gene clustering in prokaryotes frequently brings together genes with common functions, we sought to determine the extent to which ubiquitylation and ESCRT related genes are found co-located within specific regions of Asgard genomes (Figure 1B and C). As previously described^58^ within a single Odinarchaeota genome the full set of putative gene-products with homology to *Ub-ESCRT* were found together within a single gene cluster (Figure 1C). We extended this analysis by developing a simple metric of gene clustering, which was applied to Heimdall-, Loki-, Thor, and the recently described Hel-archaeotal genomes, all which harbour *ESCRT* genes^57,58,67^. This was achieved by measuring the fraction of genomes within each phylum in which each pair of genes co-localises within less than 10 kb (Figure 1B; white indicates no evidence of co-location and deep purple indicates full co-location)^68^. This analysis revealed cases where the entire set of genes was clustered together in genomes Hel and Heimdall archaea genomes, and was found organised into two relatively discrete Ub and ESCRT genomic regions in Lokiarchaeota (Figure 1B). In addition, we observed a consistent pattern of association across genomes in which the genes for ESCRT-III and Vps4 were found most tightly associated with homologues of Vps25 (Figure 1B-C, S3). This is striking as Vps25 is the subunit of the ESCRT-II complex in eukaryotes that recruits ESCRT-III to membranes, triggering vesicle budding^1–18^. Vps22/Vps36 homologues (ESCRT-II components) were found to have a similar but slightly less consistent pattern of co-location with ESCRT-III and Vps4, and the gene was usually closely associated with Vps25 (Figure 1B-C, S3).

During this analysis, we also noted that, whereas eukaryotic Vps22 and Vps25 function as part of a single hetero-tetrameric complex together with a divergent Vps22 homologue, Vps36, Asgard archaeal genomes possessed a single gene encoding a Vps22/Vps36-like protein (Figure 1A and C). A phylogenetic analysis was used to confirm that the Asgard Vps22-like protein is a closer homologue of Vps22 than it is of Vps36 (Figure 1D, Figure S4). This suggests that the eukaryotic-specific protein, Vps36, arose from a Vps22/Vps36 homolog after the divergence of Vps22 and Vps25 in Asgard archaea.

Asgard genomes also code for clear homologues of the eukaryotic ESCRT-I machinery, including homologues of Vps23 (which contain a Ubiquitin E2-Variant or “UEV” domain) and Vps28, both containing steadiness box (SB) domains. Interestingly, these genes were not tightly clustered in Asgard genomes (Figure 1B-C, Figure S5). Furthermore, the organisation of this set of genes was variable across lineages; including genomes in which individual domains were brought together to form fusion proteins (Figure 1B-C, Figure S5-8). In addition, Asgard archaea were found to lack a clear homologue of the eukaryotic Vps37. This ESCRT-I subunit is a homologue of Vps23 (reference 68 and Figure S5-8), implying that Vps37 may have arisen later in evolution in the branch leading to eukaryotes. In summary, with the notable exception of the Thorarchaeota, which seemingly lack true ubiquitin homologues (Figure 1A-C and Figure S5), most Asgard genomes have the potential to encode proteins that together resembles large parts of the conserved eukaryotic Ub-ESCRT system.

### Asgard archaeal ESCRT-I subcomplexes bind ubiquitin

In eukaryotes, a variant of the ubiquitin E2 (UEV) domain plays a key role in Ub recognition by ESCRT-I^69^. The UEV domain is similar in structure to the E2 region of the ubiquitin conjugating E2 enzymes, but lacks the key catalytic cysteine. We identified a number of similar proteins in Asgard archaea. The Heimdallarchaeota AB125 genome codes for four E2-like candidates: HeimAB125_07740, HeimAB125_09840, HeimAB125_11700 and HeimAB125_14070 (Figure S7A-B). Structural models of these Heimdall proteins were generated using I-TASSER^70–72^ (Figure S7A) and structural superimposition used to confirm that, while two of the Heimdall proteins (HeimAB125_07740 and HeimAB125_09840) possess putative catalytic Cys residues (characteristic of *bona fide* E2 ubiquitin-conjugating proteins), two (HeimAB125_11700 and HeimAB125_14070) did not contain cysteine residues at the expected catalytic positions (can we HIGHLIGHT this cysteine in Figure S6? – its hard to find!), raising the possibility that these may have UEV domains and function as ubiquitin-binding proteins (Figure S7A and B). One of these UEV domain-containing proteins represents a fusion of a Vps28 domain to UEV-Vps23 (containing ESCRT-I signatures: antiparallel coiled-coil stalk region and a steadiness box). This organisation raises the possibility that this Heimdallarchaeota protein harbours both Vps23-like ubiquitin-binding activity (via the UEV-domain) and Vps28-like functions, which are central to ESCRT-II subcomplex recruitment (Figure 2A-C and Figure S7 and S8). A phylogenetic analysis was used to confirm that both Asgard and eukaryotic UEV-like proteins cluster with eukaryotic Vps23 homologues and away from *bona fide* Ubiquitin E2 enzymes – suggesting a divergence in the structure and function of UEV and E2 domains that predates the Last Asgard and Eukaryotic Common Ancestor (LAsECA) (Figure 1E). Other Asgard species, most notably Odinarchaeota archaeon LCB_4, possess separable and distinct Vps23-like (containing the UEV domain) and Vps28 proteins, as seen in eukaryotes (Figure S8). Both these Odin proteins possess steadiness boxes, which in eukaryotes are critical for the assembly of the ESCRT-I subcomplexes^9,73,74^.

**Figure 2.**
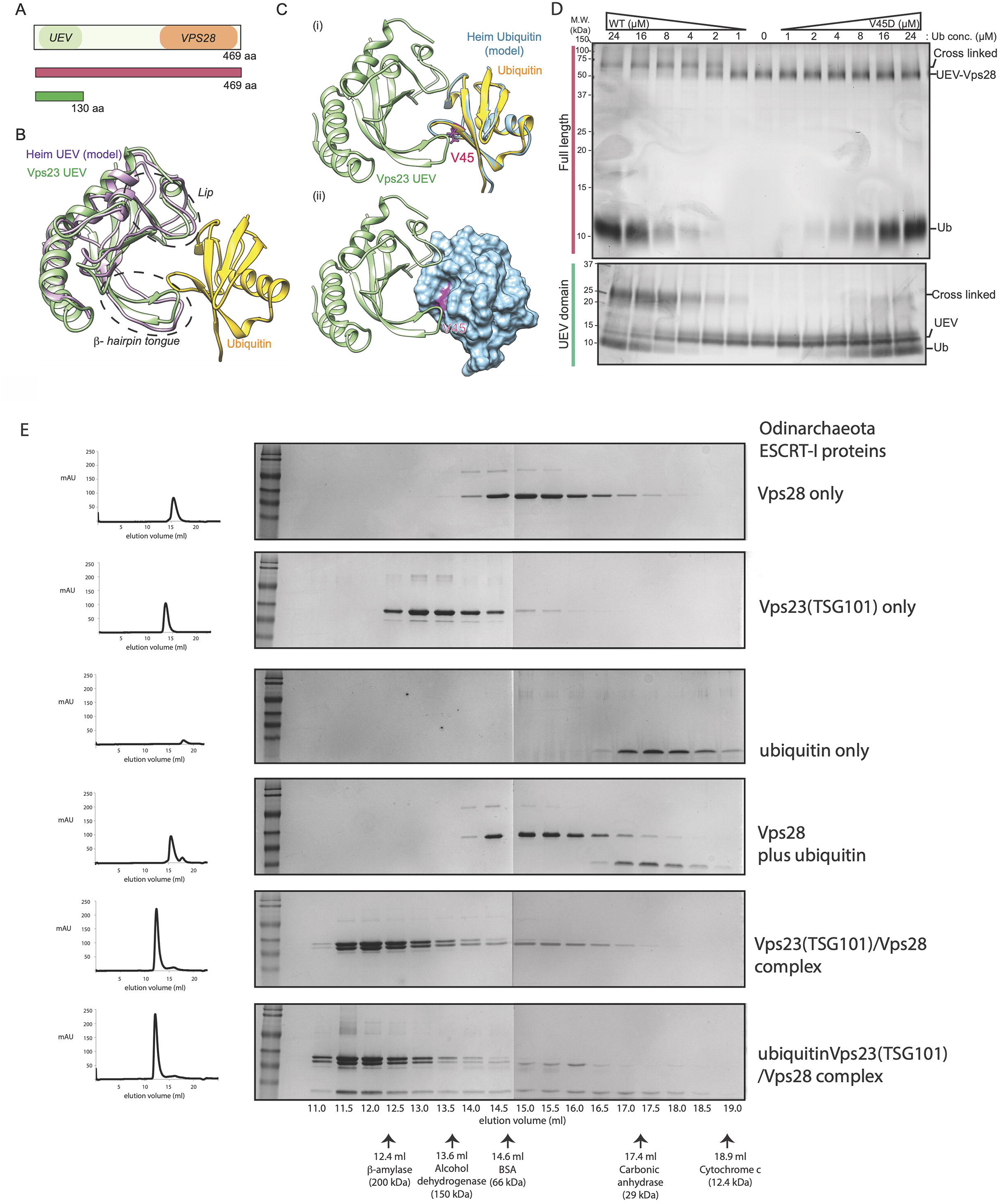
Assembly of Asgard ESCRT-I complexes with ubiquitin binding to the UEV domain of Vps23(TSG101). Schematic diagram of the domain structure of HeimAB125 Vps28 and the truncation design used in the experiments. The N-terminal ubiquitin E2 variant domain (“UEV”) and the core domain of Vps28 (“Vps28”) identified previously are highlighted. A model of the three-dimensional structure of HeimAB125 UEV. The three-dimensional structure model was created by templating the structure of the budding yeast Vps23 bound to ubiquitin. Vps23 (PDB: 1UZXa, light green) and HeimAB125 UEV (model structure, purple) are superimposed. Ubiquitin bound to Vps23 (PDB: 1UZXb) is shown in yellow. Two ubiquitin-binding “arms” of Vps23, “beta-hairpin Tongue” and “Lip” are highlighted by circles with dashed lines. The three-dimensional structure model of HeimAB125 Ubiquitin was created and superimposed with the structure of Ubiquitin in complex with Vps23 (PDB: 1UZXb). Ubiquitin (PDB: 1UZXb, yellow), model structure of HeimAB125 Ubiquitin (light blue), Vps23 (PDB: 1UZXa, light green) were shown. An amino acid residue (Val45) on the model structure located in the Ubiquitin hydrophobic patch, which is important for Ubiquitin-UEV interactions, is highlighted in magenta. The structure of HeimAB125 ubiquitin is illustrated in the ribbon diagram (i) and the surface model (ii). HeimAB125 UEV binds ubiquitin in a manner dependent on a hydrophobic patch. The interaction between HeimAB125 Ubiquitin (wild-type or V45D mutant) and UEV-Vps28 (full-length, top panel) or UEV domain (bottom panel) was tested by BS3-mediated chemical cross-linking, followed by SDS-PAGE to detect the changes in their molecular weight. Size-exclusion chromatography analysis of the Odinarchaeota ESCRT-I subcomplex assembly. Physical interaction between the *Odinarchaeota* ESCRT-I subcomplex and ubiquitin as demonstrated by size exclusion chromatography. From Top to Bottom: Vps28 protein only (top); Vps23(TSG101) protein only; ubiquitin only; Vps28 pre-incubated with ubiquitin (no interaction); Vps23 (TSG101) pre-incubated with Vps28 (stable complex formation); Vps23 (TSG101) pre-incubated with Vps28 and ubiquitin (bottom – ubiquitin binds to the Vps23(TSG1010)/Vps28 complex, via the UEV domain of Vps23(TSG101). For additional controls see Supplementary Figure S10). All proteins were separated on a Superdex S200 HR 10/300 size exclusion chromatography column. The relative elution volumes of the size standards β-amylase (200 kDa), alcohol dehydrogenase (150 kDa), bovine serum albumin (BSA) (66 kDa) and carbonic anhydrase (29 kDa) and cytochrome-c (12.4 kDa) are also indicated (in grey). Eluted fractions were resolved by SDS-PAGE and visualised by Coomassie stain. Left: chromatography UV traces (at 280 nm) for the respective elution profiles.

To test for physical interactions between these putative ESCRT-I proteins, we recombinantly expressed in *E. coli* and purified Heimdallarchaeota and Odinarchaeota ubiquitin together with their corresponding putative UEV-containing proteins and performed *in vitro* binding experiments. In the case of the Heimdallarchaeota proteins, the interaction between the purified full-length UEV-Vps23-Vps28 fusion protein and ubiquitin was analysed by chemical cross-linking followed by SDS-PAGE (Figure 2D). This revealed an increase in the apparent molecular weight of the protein upon ubiquitin binding and cross-linking (Figure 2D, top panel). In addition, we observed a ubiquitin-dependent increase in molecular weight for the isolated Heimdall UEV (Figure 2A, green), indicative of direct ubiquitin binding by this domain, rather than a different part of the full-length Vps23-Vps28 protein (Figure 2D, bottom panel). In eukaryotes, binding is mediated by an Ile-residue at position 44 (I44) of Ubiquitin, which is part of a hydrophobic patch^13^. A model of Heimdall ubiquitin was superimposed on an available crystal structure of ubiquitin in a complex with UEV (Figure 2B-C) to identify the equivalent residues (V45 in Heimdall ubiquitin). When tested experimentally, the V45D mutation dramatically reduced the interaction between ubiquitin and the UEV domain (Figure 2D) – implying that this interaction resembles the one seen in eukaryotes.

The same was true of the equivalent Odinarchaeotal proteins. When Odin homologues of ubiquitin, Vps23 and Vps28 were purified and mixed *in vitro*, we observed the formation of a stable complex by size-exclusion chromatography (Figure 2E); mediated by an interaction between ubiquitin and the UEV domain-containing Vps23 (Figure 2E, S9, and S10A). Furthermore, when analysed by size-exclusion chromatography, Vps23 migrated through the column faster than expected for a monomer (Figure S9). Follow up size-exclusion chromatography coupled with multi-angle light scattering (SEC-MALS) analyses (Figure S10 A-C) revealed that the Vps23 protein forms a stable dimeric assembly (60.37 kDa), while the calculated mass of the Vps23-Vps28 complex was consistent with a single subunit of Vps28 associating with the Vps23 dimer (yielding a combined molecular weight of 88.27 kDa). In this, the Odinarchaeotal Vps23-Vps28 complex appears like the eukaryotic ESCRT-I subcomplex, which assembles into a heterotrimeric core ‘headpiece’ comprised of Vps23, Vps28 and Vps37^74–76^. As Asgard genomes encode Vps23-like proteins but lack their Vps37 homologues (Figure 1A), it is likely that the eukaryotic ESCRT-I (Fig. S8) arose from a heterotrimeric archaeal ESCRT-I subcomplex composed of a Vps23 homodimer and Vps28.

### Potential gene duplication and functional divergence of the eukaryotic ESCRT-II subcomplex from putative Asgard archaeal precursors

Turning to the ESCRT-II subcomplexes, our genomic analysis identified close homologues of Vps22 and Vps25 in Asgard archaeal genome assemblies, but not full-length Vps36 homologues. Since eukaryotic Vps22 and Vps36 are structurally related^77,78^, and form a Vps22-Vps36 heterodimer, we investigated whether Asgard Vps22 homologues might homodimerize. To test this idea, we used size exclusion chromatography (Figure 3A) and SEC-MALS (Figure 3B) to show that a Heimdall-Vps22 homologue (HeimAB125_14050) migrates with a size consistent with it forming a dimer. Moreover, after chemical crosslinking to stabilize the Vps22 complex, through the addition of BS3 (a homo-bifunctional cross-linker) or EDC (hetero-bifunctional cross-linker that generates zero-length isopeptide bonds), the Vps22 protein band detected using SDS-PAGE had a molecular weight of about 50 kDa, corresponding to a dimer (Figure 3C). Finally, using chemical cross-linking coupled with mass spectrometry (XL-MS), we identified two dimerization surfaces (41-47 and 160-167 amino acid regions) in the HeimAB125_14050 Vps22 homodimer (Figure 3D-E and Figure S11). Taken together, these data strongly suggest that Heimdallarcheota AB125 Vps22 forms a homodimer. Thus, it appears likely that the eukaryotic Vps22-Vps36 heterodimer arose during eukaryotic evolution following a gene duplication and diversification event – just as seems to have been the case for the ESCRT-I complex. It is notable, however, that when a SEC-MALS investigation was performed with the Odinarchaeota Vps22, we did not find evidence for its homodimerization. Since the Odin protein was observed to be monomeric (Figure S10D), it is currently unclear if it adopts a homodimeric architecture in the native host conditions or might instead form a heterodimer by interacting with as-yet undiscovered proteins.

**Figure 3.**
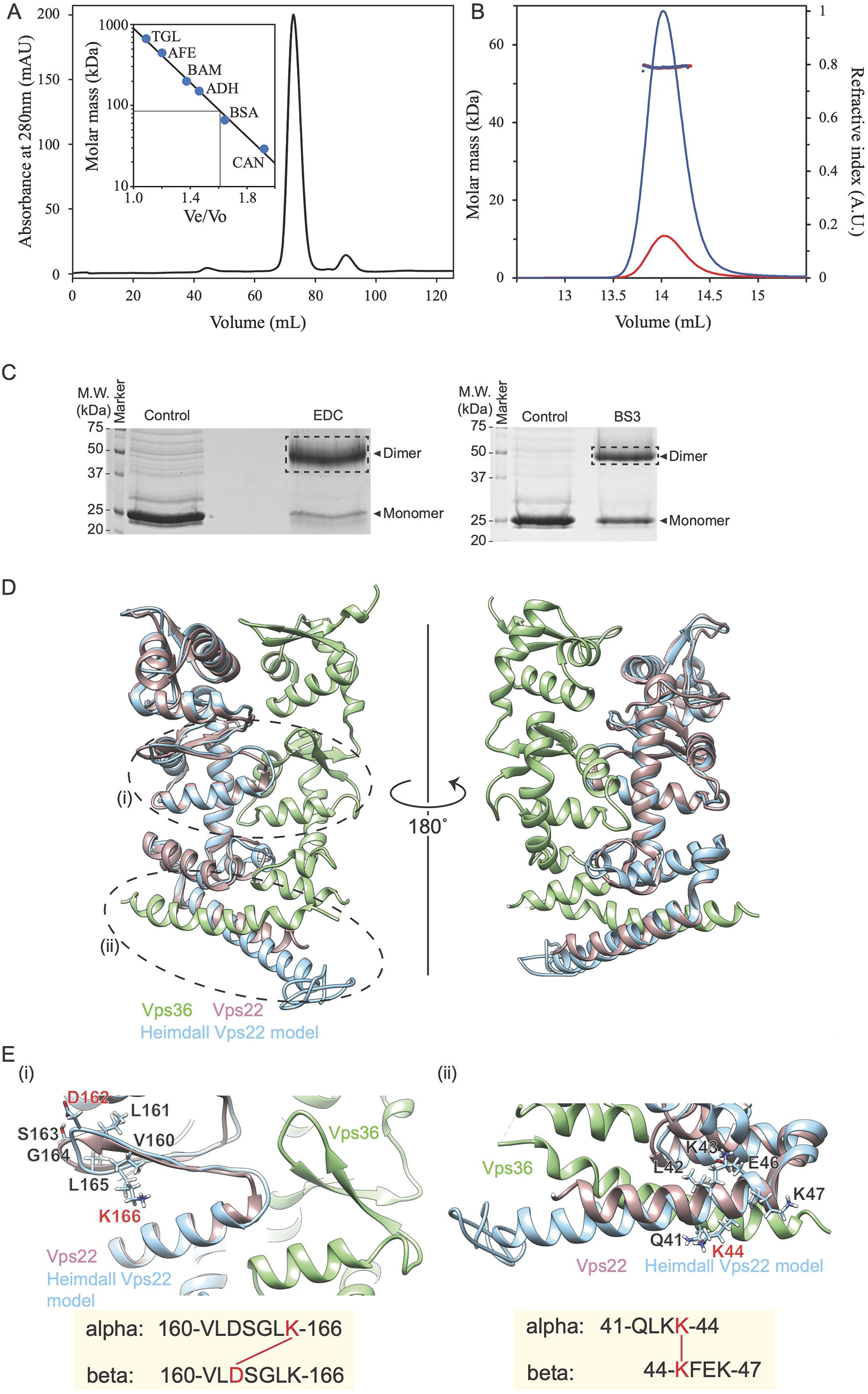
Heimdall Vps22 forms stable dimers. Elution chromatogram of Vps22 (27.9 kDa) using a Superdex 200 16/600 size exclusion column. The inset shows the column calibration curve established with standard proteins (see Methods section). Grey lines indicate the Ve/Vo and predicted molar mass (85 kDa) of Vps22. This assay suggests that this protein forms a trimer or an elongated dimer. SEC-MALS analysis of Heimdall Vps22 using a Superdex 200 increase 10/300 analytical column. The chromatograms display the calculated molar mass of the peaks (kDa) and refractive indexes (A.U.) as dots and lines, respectively, for loaded sample concentrations of 2.0 (blue) and 0.5 (red) mg/ml. The estimated masses are 54.4 and 54.2 kDa for the two protein concentrations, indicating stable formation of a Vps22 dimer, as the theoretical dimer mass is 55.9 kDa. Purified HeimAB125 Vps22 showed slower migration on SDS-PAGE gel by chemical crosslinking, whose mobility is consistent with that of a cross-linked dimer. (D and E) A model structure of HeimAB125 Vps22 superimposed on Vps22 in the structure of ESCRT-II complex (3CUQ). Side chains of dimerized peptide around 160:166 aa (ii) and 41-47 aa (ii) on the Vps22 model structure were shown in the panel. The cross-linked positions on the peptide sequence were highlighted in red.

Structural models for the Odinarchaeotal and Hemidallarchaeotal ESCRT-II subcomplex proteins using the I-TASSER^70–72^ and tr-Rosetta^79^ servers (Figure S12 A-B) suggested them having structural similarities with their eukaryotic counterparts, Vps22 and Vps25, based largely on the possession of shared tandem winged-helix (WH) domains^77,78,80^. We examined the recombinant Odinarchaeota Vps22 and Vps25 by CD spectroscopy (Figure S12 C-F). This confirmed that both proteins are folded and have the predicted secondary structural elements. To further characterise Asgard ESCRT-II subunits, these proteins were submitted to crystallisation trials. In the case of Odinarchaeota Vps25, while we were unable to crystalise the full-length soluble protein, high quality crystals could be generated using an N-terminally truncated version (deleting N-terminal residues 1 to 58), which were then used to determine the structure at a resolution of 1.80 Å (Supplementary Table 1 and Figure 4A-C). Inspection of this structure revealed that the Asgard Vps25 core is composed of a tandem WH domain repeat, consistent with the predicted tr-Rosetta model, and is very similar in structure to the equivalent eukaryotic protein (Figure 4C). Taken together, these findings provide further support for the idea that all the ESCRT-II subcomplex proteins in eukaryotes and Asgard archaea alike share a common molecular architecture based on a core tandem WH domain^77,78^. These data reinforce the concept that the ESCRT-II complex arose during archaeal and eukaryotic evolution through a series of gene duplication and specialization events^81,82^.

**Figure 4.**
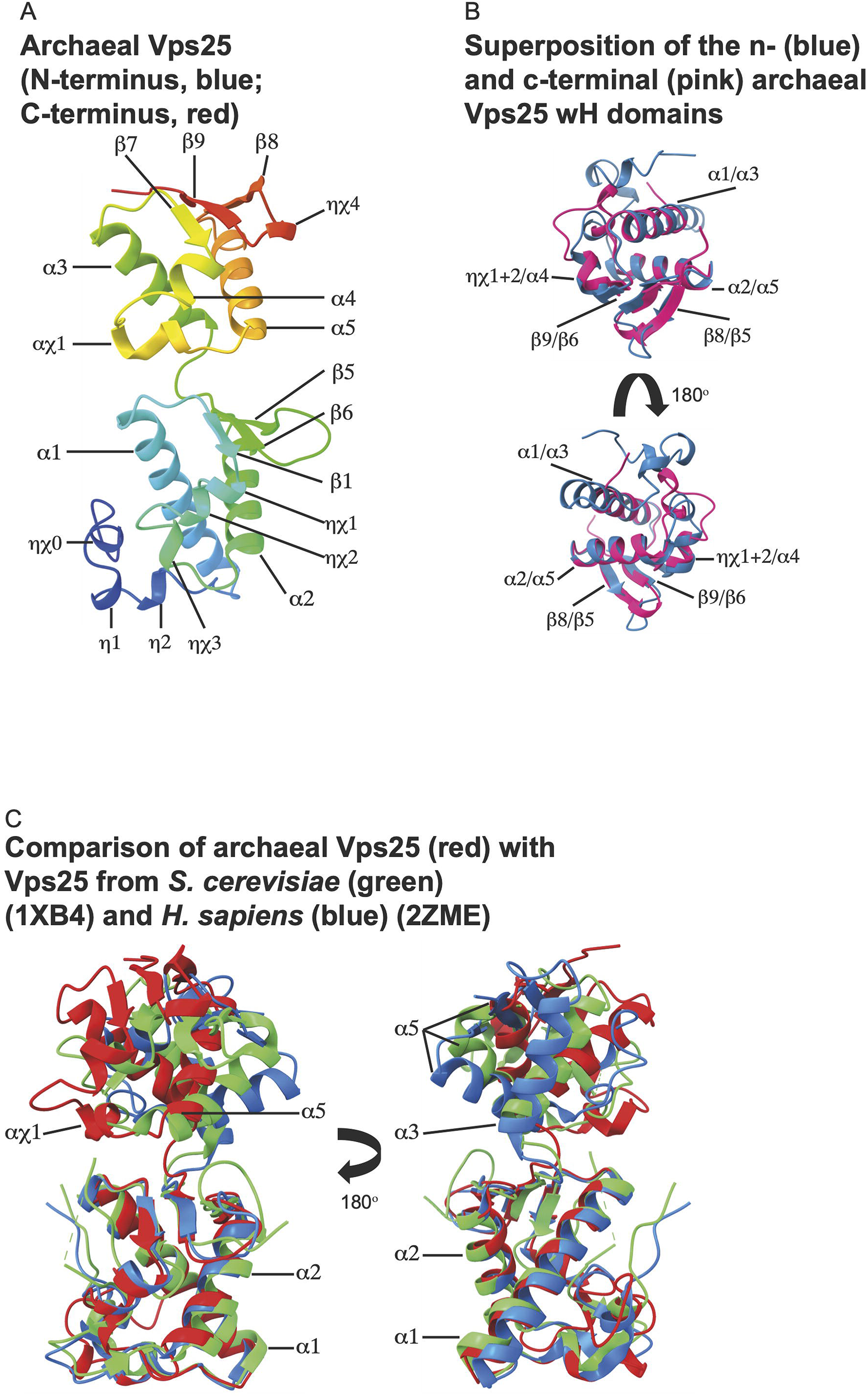
Crystal structure of the Odinarchaeota Vps25ΔN tandem winged helix (WH) domain. Archaeal Vps25ΔN tandem WH domain structure coloured from blue to red (N-terminus to C-terminus) shown in ribbon form, with secondary structural sequence elements indicated. Structural alignment of the Odinarchaeota Vps25ΔN (red) with Vps25 from *S. cerevisiae* (green) (PDB: 1XB4) and *H. sapiens* (blue) (PDB: 2ZME). Superposition of the N-(blue) and C-terminal (pink) archaeal Vps25 WH domains. Refinement and model statistics are shown in Supplementary Table 1.

### Key ESCRT complex protein-protein interactions revealed by yeast-2-hybrid analyses of the Asgard archaeal systems

Since the ESCRT-I and ESCRT-II systems characterised above function as potential protein bridges that physically connect Ub-modified proteins with the ESCRT-III machinery, we wanted a systematic way to sensitively test for interactions within and across different subcomplexes. To do so, we carried out a comprehensive reciprocal yeast two hybrid analysis (Y2H) in budding yeast to identify pairwise protein interactions for the full set of Ub-ESCRT homologues from Thor-, Odin-, Loki- and Heimdallarchaeota (Figure 5A). As a control for this analysis, we used the same approach to systematically probe for interactions between proteins that are known to function as part of the ESCRT system in the fission yeast *Schizosaccharomyces pombe* (Figure S13 and Figure S14). Importantly, in these control experiments we were able to identify many of the expected interactions between components of the Ub-ESCRT system in fission yeast. This included the previously reported interactions within the respective ESCRT complexes; ESCRT-I (Sst6-Vps28), ESCRT-II (Vps22-Vps25, Vps36-Vps25), ESCRT-III (Vps20-Vps32, Vps24-Did4), as well as published interactions that bridge the eukaryotic ESCRT-I and −II subcomplexes (Vps28-Vps36) and those that connect ESCRT-II and −III (Vps20-Vps25, Vps20-Vps22) (Figure S13A).

**Figure 5.**
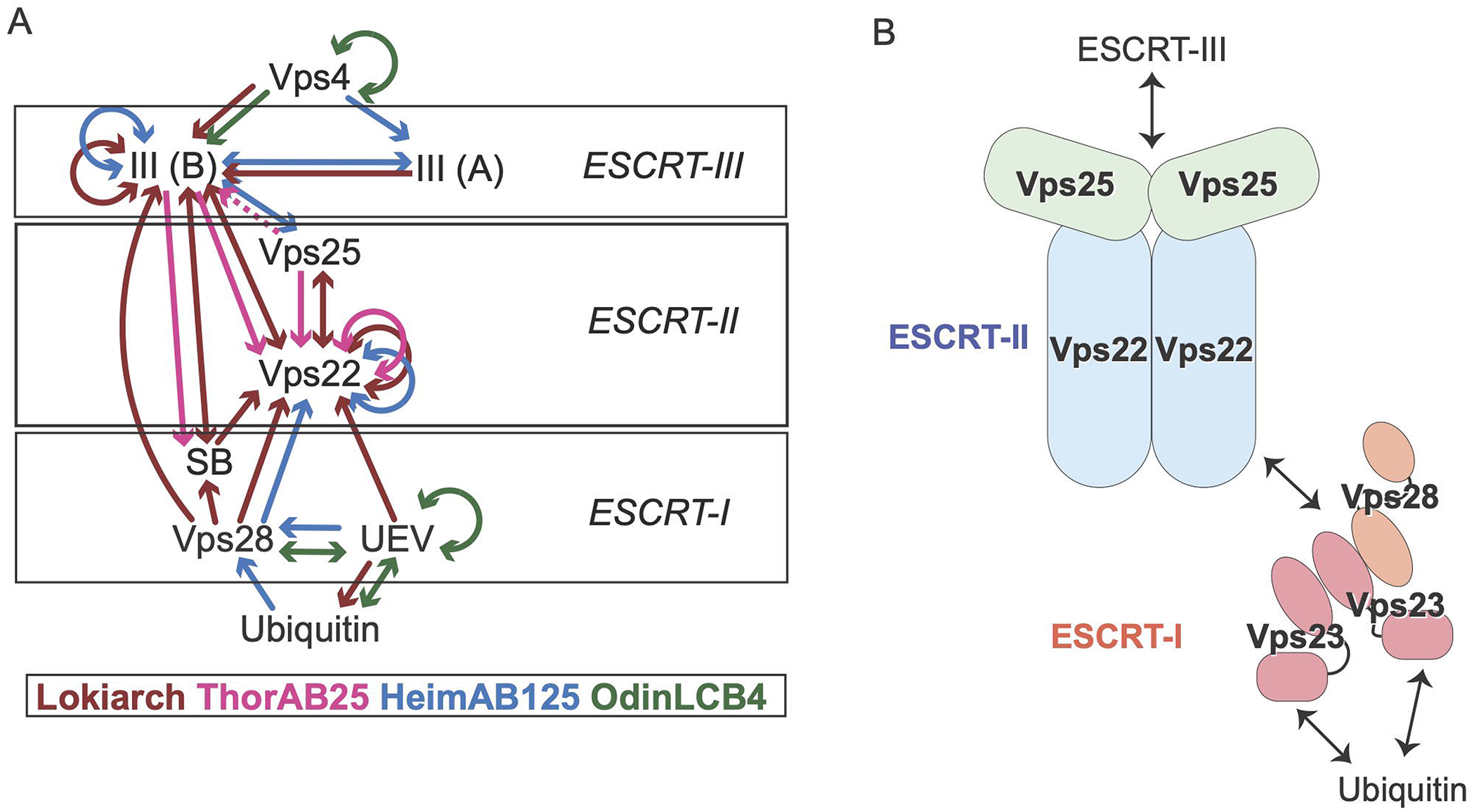
Systematic reciprocal Yeast two-hybrid assays between Asgard ESCRT proteins and new insight gained from investigating ESCRT from Asgard. Summary of Y2H interactions. Molecules related to the Ub-ESCRT pathway found in Lokiarchaeota (Lokiarch), Heimdallarchaeota (HeimAB125), Thorarchaeota, and Odinachaeota (OdinLCB4) were examined comprehensively using Y2H, and the detected interactions are illustrated (see also Figure S6 for the individual results) Schematic representation of the arrangement of the Asgard ESCRT pathway based on this work. In the Heimdallarcheota and Lokiarchaeota ESCRT-II complexes the Vps22 subunit forms a homodimer comparible to the eukaryotic Vps22/Vps36 ESCRT-II heterodimeric stalk. In Odinarcheota, however, the Vps22 homologue does not appear to dimerize and as yet undetermined factor therefore likely bridges the interaction between the ESCRT-I and −II subcomplexes. The Odinarcheota Vps23 homologue forms a dimer thereby presenting two ubiquitin-binding UEV domains. The Vps23 dimer interacts with a single Vps28 protein thus forming a tripartite complex, reminiscent of the eukaryotic Vps37/Vps23/Vps28 complex. In Heimdallarcheota the Vps23 and Vps28 functions are fused in a single protein that also dimerises. Compare with the eukaryotic arrangement as shown in Supplementary Figure S1.

We then applied Y2H analyses to systematically search for protein-protein interactions between ESCRT-related components encoded by Asgard archaeal genomes (Figure 5A). As expected, these assays revealed interactions between ESCRT-III components and the Vps4 ATPase. In line with the biochemical data presented above, these Y2H analyses also identified interactions between ubiquitin and UEV domain-containing proteins in Heimdallarchaeota, Odinarchaeota and Lokiarchaeota (Figure 5A and S14). Furthermore, the UEV domain-containing Vps23 homologues from Odinarchaeota and Heimdallarchaeota displayed interactions with Vps28, as expected if they formed an ESCRT-I subcomplex, whereas the Heimdallarchaeota utilise the single Vps23-Vps28 fusion protein homologue discussed above. The Y2H assays also detected the interaction between ubiquitin and this Heimdall ESCRT-I fusion (Figure 5A and Figure S13B). Interestingly, this analysis also suggested an alternative pattern of protein interactions for the corresponding ESCRT-I proteins from Loki and Thor. In these two cases, the freestanding steadiness box protein, equivalent to the alpha-helical hairpin ‘headpiece’^73,74,83^ coded within the same genomic neighbourhood as the rest of the Loki ESCRT machinery (Figure 1C), may come into play by interacting with Vps28 – mirroring the role of its eukaryotic counterpart in mediating interactions between Vps23, Vps28 and Vps37.

The Y2H analyses also identified multiple interactions between the Lokiarchaeota and Heimdallarchaeota ESCRT-I (Vps28 and/or Vps23) and ESCRT-II (Vps22) complexes. In addition, the Y2H experiments indicate that the ESCRT-II component Vps22 from Heimdall, Loki and Thor can interact with themselves, in agreement with the biochemical assays shown above. Finally, we identified numerous interactions linking the Asgard ESCRT-II and −III (Vps25-ESCRT-IIIB), Vps4-ESCRT-III and between the ESCRT-III homologues (-IIIA and −IIIB) (Figure 5A and Figure S14), and suggests the possibility of additional interactions between ESCRT-I and II subcomplexes with ESCRT-III proteins that bypass Vps25 in Loki and Thor archaeota. We note that although we were unable to detect many such interactions between the Odinarchaeotal proteins using the Y2H approach, since Odin is a thermophile, it seems likely that the temperature used for these experiments (25°C) may have influenced our ability to identify interactions between proteins that are optimised to fold and work at much higher physiological temperatures^58^.

## DISCUSSION

Here we provide experimental and computational support for the idea that many Asgard archaea possess a streamlined version of the Ub-ESCRT system present in eukaryotes (Figure 5B). It is clear from our analyses, however, that the precise composition differs between species and across phyla. This is especially the case for ESCRT-I subunit architecture. However, except for Thorarchaeota (in which ubiquitin encoding genes are yet to be identified), we found ubiquitin-binding UEV-domain containing proteins coded by the genomes analysed. While these domains were often harboured within proteins homologous to Vps23, which include a C-terminal alpha-helical headpiece region involved in ESCRT-I complex assembly, in other systems this alpha-helical ‘steadiness box’ domain was encoded by a freestanding protein. Alternative ESCRT-I domain arrangements were also observed, such as the UEV-Vps23-Vps28 fusion found in Heimdallarchaeota (Figure S8). Taken together, the clear synteny between genes of ubiquitylation apparatus and the ESCRT machinery in the Loki-, Hel-, Odin-, and Heimdallarchaeotal genomes, and the experimentally verified Y2H interactions between ubiquitin and UEVs in all the Asgards investigated, support a model in which ubiquitylated substrates recruit the ESCRT-I in Asgard archaea.

In line with this, we were able to demonstrate direct binding between ubiquitin and UEV-containing proteins that was dependent on a conserved hydrophobic patch in ubiquitin. Although Asgard archaea appear to lack homologues of the eukaryotic ESCRT-I subunit Vps37 (which assembles into a trimer with Vps23 and Vps28), it is notable that Vps23 and Vps28 from Odinarchaeota assemble into a similar trimer that contains two copies of Vps23. It is therefore possible that all three proteins of the eukaryotic ESCRT-I complex evolved from a common ancestor containing an alpha-helical hairpin region, with Vps37 having arisen as a eukaryotic innovation^73^.

Similarly, the eukaryotic ESCRT-II subcomplex forms a ‘Y-shaped’ hetero-tetrameric structure consisting of a Vps22/Vps36 stalk which binds two Vps25 subunits^77,78^. Although Vps22 and Vps25 coding genes were readily identifiable in the Asgard genomes as reported previously, we were unable to identify Vps36 homologues. However, our biochemical and Y2H interrogation suggest that Vps22 from several Asgard phyla likely forms homodimers. In the corresponding eukaryotic complex, all three of these ESCRT-II proteins (Vps22, Vps36 and Vps25) contain an evolutionarily conserved globular core consisting of tandem winged helix (WH) domains^77,78,80^. The same appears true for the Asgard ESCRT-II machinery, based upon modelling of Vps22 and Vps25 proteins from Odinarchaeota, and the crystallographic Odinarchaeotal Vps25 structure. This suggests that all ESCRT-II proteins were initially derived from a single WH-domain protein progenitor, with Vps36 emerging during eukaryogenesis.

Do ESCRT-I, −II, and −III contribute to a single pathway? The colocation and synteny analyses strongly suggest this possibility. The colocation of ESCRT-III with Vps25 indicates that these proteins function in a related biochemical process. Y2H experiments revealed consistent interactions between Vps25 and ESCRT-IIIB. Furthermore, this analysis provided support for there being an interaction between ESCRT-I and ESCRT-II. We also found an interaction between the Lokiarchaeota steadiness box protein and Vps22, supporting the existence of physical interactions between the ESCRT-I and ESCRT-II subcomplexes. These Y2H data and the corresponding gene cluster analysis point to ESCRT −I, −II, and −III functioning in concert in Asgard archaea – although confirmation of this will require a future cell biological and/or biochemical analyses. While it is not yet clear how the Ub-ESCRT system evolved, we note that while ESCRT-III type proteins can be traced back to the last universal common ancestor (LUCA)^63^, the Vps4 and ESCRT-III pair can only be found in archaea, whereas Ub, ESCRT-I and ESCRT-II components are only found together in Asgard (with the exception of Thorarchaeota, which seemingly lack true ubiquitin homologues). This suggests a plausible pathway for the stepwise evolution of the eukaryotic ubiquitin-directed ESCRT-dependent membrane trafficking system as, from its simple beginnings in an archaeal progenitor, the machinery grew in complexity through successive rounds of domain concatenation, gene duplication and divergence (Figure 5B).

## MATERIALS AND METHODS

### Genomic survey of protein homologues

All genomes from organisms classified as Asgard archaea were downloaded from NCBI on December 5^th^, 2020. These genomes were taxonomically reclassified through a phylogenetic analysis based on a set of 15 ribosomal proteins encoded in co-locating genes^84^. To ensure annotation homogeneity, protein sequences were predicted de novo using Prodigal v2.6.3^85^, and ribosomal protein genes were detected using psiblast^86^ using predetermined orthologous sequences^58^, aligned with Mafft-linsi v7.450^87^ and processed with trimAl v1.4.rev22^88^ to remove sites with over 50% gaps. All genomes containing at least 5 of these proteins were concatenated and used to reconstruct a tree with IQ-Tree v2.0-rc1^89^ the LG+C60+R4+F model, using 1000 pseudoreplicates for ultrafast bootstrap^90^ and SH-approximate likelihood ratio tests (Figure S1).

These reclassified genomes were then annotated using interproscan v5.48-83.0^91^. A set of interpro domains were used as diagnostic for the inference of Vps23/37 (IPR017916), Vps28 (IPR007143, IPR037206), Vps23/37/28 (IPR037202), Vps22/36 (IPR016689, IPR040608, IPR021648), Vps25 (IPR008570, IPR014041), ESCRT-III (IPR005024), Vps4 (IPR007330, IPR031255, IPR015415), ubiquitin (IPR029071, IPR000626), ubiquitin-activating enzyme E1 (IPR000594), ubiquitin-conjugating enzyme E2 (IPR000608, IPR006575, IPR016135), ubiquitin-ligase enzyme E3 (IPR018611), and deubiquitinating enzyme (IPR000555) genes.

Co-location of these genes was investigated through custom perl scripts and visualised using R (R Core Team 2018, R: A language and environment for statistical computing. R Foundation for Statistical Computing, Vienna, Austria. https://www.R-project.org/) and the packages ggplot2^92^, cowplot (https://CRAN.R-project.org/package=cowplot), and genoPlotR^93^.

### Generation of model structures using I-TASSER server

The model structures used in this study were generated as described below:

1. The model structure of Heimdall E2-like proteins for Figure S7A were generated using I-TASSER^70–72^ (https://zhanglab.ccmb.med.umich.edu/I-TASSER/). No template structure was selected for this modelling. The amino acid sequence used for structural modelling is as follows; HeimAB125_07740 (27th-135th amino acid residues); (ii) HeimAB125_09840 (1st-107th amino acid residues); (iii) HeimAB125_14070 (51st-108th amino acid residues); (iv) HeimAB125_11700 (25th-122nd amino acid residues). The amino acid residues for the structural modelling were selected based on InterPro annotation of “UBIQUITIN_CONJUGAT_2”. It is important to note that the c score of HeimAB125_14070 truncation was NaN, due to lower reliability of the model structure.
2. The model structure of Heimdall UEV domain in HeimAB125_14070 (1st-130th amino acid residues) in complex with ubiquitin (used in Figure 3B and S7B) was generated using I-TASSER with crystal structure of yeast UEV in complex with ubiquitin (PDB: 1UZXa).
3. The model structure of Heimdall UEV domain in HeimAB125_14070 (1st-130th amino acid residues) for Figure S7A was generated by using I-TASSER with crystal structure of UEV (PDB: 1KPQ).
4. The model structure of Heimdall ubiquitin (HeimAB125_14240) and Heimdall Vps22 (HeimAB125_14050) were generated by I-TASSER without assigning template structures.

In most of the cases, I-TASSER provided several models. We always used the model which showed the highest c scores. The score, templates and amino acid region were listed in Table S1.

### Asgard proteins used in this study

The Heimdall-, Loki-, Odin-, and Thorarchaeota amino acid sequences were obtained from Uniprot (https://www.uniprot.org/) and the Uniprot Entry ID are listed in Table S2. The corresponding genes were synthesized for the expression in *E. coli* and yeast. The Odinarchaeota_LCB4 ORFs (accession codes provided in Supplementary Table 2) were PCR amplified form MDA amplified environmental DNA isolated from the Lower Culex Basin Yellowstone National Park, USA as described^94^ and cloned into either pET28a (Vps23 and Vps28) or pET30 (Vps22 and Vps25) (Novagen), respectively. Amplified genes were cloned into the plasmids using the restriction sites placing the ORFs in frame with the plasmid-encoded hexa-histidine tags.

### Plasmids used in this study

The Asgard genes obtained by gene synthesis were cloned into yeast two-hybrid (Y2H) vectors and *E. coli* expression vector. The plasmids for Y2H are listed in Table S3. The *E. coli* expression plasmids are listed Table S6.

### Systematic, Reciprocal Yeast Two-Hybrid assays

Y2H assays were performed using the set of genes listed in Table S3. The plasmids used in this study are listed in Table S3. Indicated genes of interest were cloned both in “bait-ProteinA” and “prey-ProteinB” vectors or vice versa, which have DNA binding protein LexA and/or activation domain of Gal4p were cloned into pMM5 and pMM6 plasmids respectively^95,96^. Plasmids carrying these constructs were transformed into the yeast strains SGY37 (MATa) and YPH500 (MATα). Transformants with plasmids plexADBD (pMM5) and pGal4AD (pMM6) were selected on plates lacking Histidine or Leucine, respectively. After mating, the two strains carrying the desired plasmids were grown on YPD plates for 2 days at 30°C and replica plated on selection plates (without Histidine and Leucine) for 2 days at 30°C before the overlay. The interaction between the protein products fused to the DNA binding and activation domains were analyzed by the activity of β-galactosidase by the cleavage of X-Gal (BIO-37035, Bioline, UK). For detecting the β-galactosidase activity overlaying of low melting agarose with X-Gal (over lay mix was prepared freshly), overlay solution was added slowly on to the plates. Interaction of LexA-Protein-A with Gal4-Protein-B resulted in the activation of expression of the lacZ gene coding for β-galactosidase, converting X-Gal to produce blue colour. Plates were monitored every 30 minutes to see the appearance of blue colour. Plates were scanned after 16hr of incubation with the X-Gal overlay mixture.

### Phylogenetic reconstruction

#### UEV and E2

Amino acid sequences of UEV domain-containing proteins, TSG101/Vps23 and UBC domain-containing proteins in *H. sapiens*, *S. cerevisiae*, *D. discoideum*, *E. histolica*, *A. thaliana*, *C. marolae*, *T. brucei*, *T. pseudonana* and *T. parva* were obtained from Uniprot. Asgard E2L proteins from Odinarchaeota (strain LCB_4), Heimdallarchaeota (strains AB125, LC2 and LC3), and Lokiarchaeota (strains GC14_75 and *CR*_4) were also obtained from Uniprot. These sequences were aligned with Mafft-linsi v7.450, and the resulting multiple-sequence alignment was used as query for a Psiblast (v2.10.0+) against all Asgard archaeal genomes (see Genome survey of protein homologs). All hits with e-values lower than 1e-5 were used together with query sequences and aligned using Mafft-linsi. The resulting alignment was trimmed using trimAl v1.4.rev22, and sequences containing over 60% gaps in the trimmed alignment were removed. The obtained alignment was used for a phylogenetic reconstruction with IQ-Tree 2.0-rc2^89^, under the model Q.pfam+C20+G4+F, chosen by ModelFinder^97^ between combinations of empirical matrices (LG, WAG, JTT, and Q.pfam) with mixture models (C20, C40, and C60) and various rate heterogeneity (none, G4 and R4) and frequency (none, and F) and using 1000 ultrafast bootstrap pseudoreplicates. The resulting phylogeny was used as guide to reconstruct another tree under the PMSF approximation of the chosen model and using 100 non-parametric bootstrap pseudoreplicates. The resulting bootstrap trees were used both using the standard Felsenstein Bootstrap Proportion and the more recent Transfer Bootstrap Expectation^98^ interpretations.

#### Vps22 and Vps36

Eukaryotic Vps22, Vps36 and Vps25 and Asgard Vps22/36 and Vps25 homologs sequences were downloaded from NCBI (Supplementary Data X). These 187 sequences were aligned using Mafft-linsi v7.450 and trimmed with trimAl with the parameter “-gappyout”. A maximum-likelihood tree was then reconstructed using IQ-Tree v2.0-rc1 under the model LG+C60+R4+F, using 1000 ultrafast bootstrap and SH-approximate likelihood ratio test pseudoreplicates. In parallel, potential outgroup sequences (eukaryotic Rpc35/Rpc6, Asgard archaeal UFM1 and bacterial ScpB; Supplementary Data X) were downloaded and added to the previous sequences. Three additional Asgard archaeal sequences were found to contain potential plekstrin domains and were used as query for a Blast-p search against the Asgard proteomes to recruit homologs identified as hits with e-values lower than 1e-10. The resulting set of 314 sequences was then aligned with Mafft-linsi v7.450 and trimmed with trimAl to remove all sites with over 90% gaps. The resulting trimmed alignment was used to reconstruct a maximum-likelihood tree using IQ-Tree v2.0-rc2 under the PMSF approximation^99^ of the LG+C60+R4+F model using 100 non-parametric bootstrap pseudoreplicates. The resulting bootstrap trees were used both using the standard Felsenstein Bootstrap Proportion and the more recent Transfer Bootstrap Expectation interpretations. To ensure we did not miss possible homologs of ESCRT-II sequences outside of Asgards, we used the previous set of 187 Vps22/Vps36/Vps25 sequences as query for a psiblast search against the NR database (1 iteration, e-value threshold of 1e-10), and parsed the resulting 4745 hits to remove proteins originating from Asgard archaeal or eukaryotic genomes. After parsing, only 9 sequences remained, belonging to various putative archaea and bacteria. A Blast-p search of these sequences against NR confirmed that their best hits were Asgard archaea or eukaryotic sequences. We added these sequences to the previous 227 Vps22/Vps36/Vps25/Outgroup sequences, aligned them with Mafft-linsi v7.450 and trimmed with trimAl to remove sites with over 50% gaps. We used this alignment to reconstruct a tree with IQ-Tree under the LG+C20+G4+F model, using 1000 ultrafast bootstrap and SH-approximate likelihood ratio test pseudoreplicates. The resulting tree confirmed that these 9 homologs were well embedded in the clades of Asgard archaeal or eukaryotic Vps22, thus likely representing Asgard archaeal or eukaryotic Vps22 sequences that have been misclassified in public databases.

### bdSUMO tag vector and bdSUMO protease used in this study

The vector carrying a 6-His residues (His-tag) followed by SUMO protein from *Brachypodium distachyon* were generated as described before^100,101^ with slight modification. The gene of *B. distachyon* SUMO protein (bdSUMO) was synthesized (IDT gBlock) and cloned into pET28a in frame with sequence encoding the N-terminal His-tag. The codon of the bdSUMO was optimized for the expression in *E. coli* K12 strain. The resulting vector, pSUMO was used to clone Heimdall ESCRT genes for their expression in *E. coli* BL21(DE3). To express and purify SUMO protease in *B. distachyo*, we synthesized genes of *B. distachyo* SENP1 whose codone were optimized for expression in *E. coli* (IDT gBlock). The gene fragment was cloned into pET28a vector in-frame with N-terminal His-tag. The protein, His-bdSENP1 was expressed in BL21(DE3) and purified and used for SUMO-TAG cleavage.

### Buffers used for protein purification

The composition of ESCRT lysis buffer was as follows: 50 mM Tris-HCl (pH 7.5), 2.5 mM MgCl2, 150 mM NaCl, 2 mM DTT, 2 mM ATP, and 15 mM Imidazole. The composition of ESCRT buffer was as follows: 50 mM Tris-HCl (pH 7.5), 2.5 mM MgCl2 and 150 mM NaCl. The composition of XL buffer is as follows; 20 mM HEPES-NaOH (pH 7.5), 150 mM NaCl.

### Protein purification

#### (A) Heimdall proteins

##### Purification of untagged proteins

The proteins were expressed as N-terminal His-SUMO fusions (His-SUMO) from a pSUMO vector. After the affinity purification using His-Nickel interaction, the His-SUMO was cleaved by His-bdSENP1 and both the N-terminal His-SUMO tag and His-bdSENP1 were absorbed on a Ni-NTA column. The untagged recombinant protein was further purified by size-exclusion chromatography (SEC).

##### Purification of ESCRT proteins

##### [A] Heimdall Vps22 and Full-length or UEV-domain of Heimdall UEV-Vps28 (HeimAB125_14070)

The E. coli cells expressing these fusion proteins were lysed using a pressure homogenizer (Stanstead #FPG12800, the 20-30 psi for several times precooled at 5°C) instead of sonicator. Elution after the SEC was concentrated followed by high-speed centrifugation at 4°C (21000xg, 15 min) to get rid of aggregation. The samples were snap-frozen and stored at −80°C.

##### [b] Ubiquitin with N-terminal His tag

Cells were re-suspended in lysis buffer containing 2x concentration of PIC (Roche complete, EDTA-free #05056489001) and 2 mM PMSF. Cells were lysed using a pressure homogenizer (Stanstead #FPG12800, the 20-30 psi for several times) precooled to 5°C. After cell lysis, the total volume of the sample was ~50 mL. The insoluble fraction was removed by high-speed centrifugation at 4°C (25658 x g for 1 hour, Thermo Fisher SCIENTIFIC #A23-6 x 100 rotor). The supernatant was incubated with 2 ml Ni-NTA resin (Thermo SCIENTIFIC #88222) for 1 hour at 4°C. The resin was washed with 200 mL ice-cold lysis buffer, followed by 150 ml ESCRT-buffer. The bound protein was eluted with ESCRT-buffer containing 300 mM imidazole. Elution fractions were combined and concentrated to a volume of 500 μl. The sample was spun at 21000 x g for 15 min at 4°C and the supernatant was applied to 16/60 sephacryl S-100 HR column equilibrated with ESCRT-buffer. The eluate was concentrated and spun at 4°C (21000xg, 15 min) to get rid of aggregates. The samples were snap-frozen and stored at −80°C.

#### [B] Odinarchaeal proteins

Thermophilic *Odinarchaeota* proteins were expressed in Rosetta (DE3) pLysS *Escherichia coli* cells (Novagen). Cultures were grown at 37°C to an OD_600_ of 0.3 then cooled to 25°C and further grown to an OD_600_ of 0.6 and induced overnight with 0.33 mM IPTG. Cells expressing the recombinant *Odinarchaeota* proteins were harvested by centrifugation, resuspended in 20 mM Tris-HCl (pH 8.0), 300 mM NaCl, 5% glycerol, 0.05% β-mercaptoethanol. 1X EDTA-free protease inhibitors (Complete cocktail, Roche) were added and cells were lysed by sonication and heat clarified at 60°C for 20 min before centrifugation at (14,000 r.p.m. for 10 min) to remove insoluble material. Supernatants were filtered and then purified by IMAC by gravity flow to a column of Ni-NTA agarose (Qiagen). The columns were washed with resuspension buffer and then resuspension buffer plus 15 mM imidazole. Proteins were then eluted in resuspension buffer plus 500 mM imidazole. Fractions containing the purified proteins were pooled and concentrated before running a size-exclusion chromatography (SEC) step over a Superdex 200 16/600 column (GE Healthcare), in 20 mM Tris-HCl pH 8, 300 mM NaCl, 5% glycerol, 0.5 mM dithiothreitol. N-terminal His-tags were then removed from the Odinarchaeota Vps23(TSG101) and Vps28 proteins by thrombin cleavage and further purification by SEC. Fractions containing the purified proteins were pooled, concentrated, aliquoted and flash frozen in liquid N_2_. Protein concentrations were quantified by UV spectrophotometry.

### Analytics Size Exclusion Chromatography

Heimdallarchaeota Vps22 (27.9 kDa) was subjected to analytical SEC using a Superdex 200 16/600 size exclusion column (GE Healthcare). The sample was loaded onto the column in a buffer comprised of 20 mM Tris-HCl pH 8.0, 200 mM NaCl and 5% (v/v) glycerol at a flow rate of 0.5 mL/min. The calibration curve was established under the same conditions using the following standard proteins (Sigma MWGF1000): carbonic anhydrase (CAN; 29 kDa), bovine serum albumin (BSA; 66 kDa), alcohol dehydrogenase (ADH; 150 kDa), beta-amylase (BAM; 200 kDa), apoferritin (AFE; 443 kDa) and thyroglobulin (TGL; 669 kDa).

Physical interaction between the *Odinarchaeota* ESCRT-I complex proteins (Vps23, Vps28 and ubiquitin) was examined by size-exclusion chromatography using an analytical Superdex S200 HR 10/300 column (GE Healthcare). Prior to the gel filtration analyses, ESCRT-I complexes were formed at 60°C by mixing 250 μg of each protein in a final volume of 500 μl gel filtration buffer (20 mM Tris [pH 8.0], 150 mM NaCl, 5% glycerol, 1 mM DTT). The complexes were subsequently spun at 16,000 *g* in a benchtop centrifuge for 5 min to remove any precipitated material, before loading onto the size exclusion chromatography column. 0.5 ml fractions were collected and resolved by SDS-PAGE, on 15% polyacrylamide gels. The proteins were then visualized with Coomassie stain.

### Size exclusion chromatography–multi-angle laser light scattering (SEC-MALS)

The molecular mass and oligomeric state of Heimdall Vps22 was determined in solution using SEC-MALS. Data were obtained with a Wyatt HeleosII18 angle light scattering machine connected to a Wyatt Optilab rEX online refractive index detector (Wyatt Technology). Samples were purified using a Superdex 200 increase 10/300 analytical gel filtration column (Cytiva) coupled to an Agilent 1200 series LC system at 0.5 ml/min in 20 mM Tris-HCl pH 8.0, 200 mM NaCl buffer before detecting the light scattering and refractive index in a standard SEC-MALS format. Protein concentration was obtained from the excess differential refractive index of 0.185 ΔRI for 1 g/ml or using the sequence UV extinction coefficient of 0.964 at 280 nm for 1 mg/ml calculated by ProtParam. The determined protein concentration and scattering intensities were used to estimate the molecular mass from the intercept of a Debye plot using Zimm’s model and the Wyatt ASTRA software. The experimental configuration was checked with a BSA standard, run in the same buffer and using the same sample injection volume of 100uL. The BSA monomer peak was utilised to examine the mass determination and to inspect the inter-detector delay volumes and band broadening parameters that were used during analysis in Wyatt’s ASTRA software. The SEC chromatogram, showing RI as concentration signal, is shown in Figure 3B as blue and red lines for loaded sample concentration of 2 and 0.5 mg/ml, respectively. The mass evaluated averaged over the central 75% of peak area is 54.4 and 54.2 kDa for the two loadings indicating stable formation of dimer at SEC concentrations. The mass evaluated using UV as concentration source was 55.7 kDa for the 2 mg/ml sample.The Odinarchaeota samples were analysed by SEC-MALS (100 μl protein complex at 2 mg/ml) were passed over a Superdex 200 10/300 Increase GL column (GE Healthcare), in 20 mM Tris (pH 8.0), 300 mM NaCl. The column output was fed into a miniDAWN TREOS MALS detector system with a 60mW laser source at 664 nm, and three fixed angle detectors at 49. 90, and 131 degrees (Wyatt Technology), followed by a Shimadzu RID-20A Refractive Index Detector at 30.5°C.

### Circular Dichroism (CD)

Proteins were buffer exchanged into freshly prepared buffer (10 mM potassium phosphate, 50 mM sodium sulphate, pH 7.2) using PD-10 desalting prepacked columns (Sephadex G-25M, GE Healthcare) following manufactures instructions.

From the protein samples (5 μM) CD spectra (in triplicate) were acquired using a Chirascan Plus Benchtop CD spectrophotometer over 180-260 nm with a bandwidth of 2 nm and a pathlength of 0.2 mm. The mean buffer subtracted CD spectra (measured ellipticity: mdeg) were interpolated between 190-250 nm using Origin Pro 2018b and fitted to the BeStSel algorithm to determine the secondary structural elements^102^. Structural models for each ESCRT protein were generated using I-TASSER^70–72^ (or for full-length Vps25, trRosetta^79^) after which the STRIDE web server^103^ was used to estimate secondary structure elements for comparison with the CD derived estimations.

### ΔN_Vps25 Crystallisation conditions

An N-terminally truncated Vps25 expression construct (removing the first 58 amino acids) was generated using the primers and OdΔN_ESCIIV25forXhoI and OdESCIIV25revXhoI as described in Supplementary Table 2. The protein was purified as described above, except 5.25 mM TCEP was used as the reducing agent in the final size exclusion chromatography step.

ΔN_Vps25 crystals were grown by sitting-drop vapour diffusion using our in-house high-throughput crystallisation platform ^104^. Vps25 was used at a concentration of 21.4 mg/ml and the best crystals were obtained in the condition E12 of the Morpheus screen^105^: [120 mM ethylene glycols, 100 mM buffer 3 (26.7 ml 1 M bicine plus 23.3 ml 1 M Trizma base), 12.5 % (w/v) PEG 3350, 12.5 % (w/v), 12.5 % (w/v) PEG 1K, 12.5 % (w/v) MPD], pH 8.5 at 20°C with a protein: reservoir ratio of 1:4 and a total volume of 0.4 μl. The condition was already cryo-protected. Crystals were harvested by flash cooling in liquid nitrogen.

### X-ray diffraction data collection

Native diffraction data were collected at Diamond Light Source (Harwell, UK) at beamline I03. Data were collected over 360° with 0.1° oscillation (Supplemental Table 1), integrated with DIALS (https://doi.org/10.1107/S2059798317017235) and scaled/merged with Aimless (https://www.ncbi.nlm.nih.gov/pmc/articles/PMC3689523/) from the CCP4 suite^106^. The crystals belong to the space group P2_1_2_1_2, with unit cell dimensions of a = 101.23 Å, b = 31.5 Å, c = 59.5 Å and one molecule per asymmetric unit. The crystals diffracted up to 1.8 Å. BALBES was used to determine initial phases by Molecular Replacement against the entire PDB^107^. Manual building was done in COOT^108^ and refinement with REFMAC5. MOLPROBITY was used for model validation^109^. Statistics are listed in Table 1. The coordinates and structure factors of the *Odinarchaeota* Vps25ΔN crystal structure were deposited in the Protein Data Bank under accession code 7PB9.

### Chemical cross-linking of proteins

#### [A] Vps22 dimer

Vps22 was diluted to 15 μM after the buffer exchange to XL-buffer and incubated with 16 mM EDC (Thermo Scientific, #22980) and 16 mM Sulfo-NHS (Thermo Fisher Scientific, #A39269) or 2 mM BS3 [bis(sulfosuccinimidyl)suberate, Creativemolecules, #001SS] on ice for 1 or 2 hours, respectively. 55.6 mM Tris-HCl (pH 6.8) was added into the mixture to quench the cross-linking reaction. The sample was incubated on ice for 10 min to quench the cross-linking reactions. The samples were loaded in SDS-PAGE gels to separate individual or cross-linked proteins.

#### [B] Ubiquitin and UEV

The full length (0.4 μM) or 1:130 amino acid residues containing UEV domain (5 μM) of Heimdall_14070 were incubated with 0, 1, 2, 4, 8, 16 and 24 μM Heimdall_14240 (wild-type or I45V) and 5 mM BS3 (Creativemolecules, #001SS) on ice for 2 hours. The cross-linking reaction was quenched by addition of 50 mM Tris-HCl (pH 6.8). The samples were loaded in SDS-PAGE gels to separate individual or cross-linked proteins. 50 mM Ammonium Bicarbonate (#A6141, Sigma) was used to dilute the sample using equal volume and reduced/alkylated using 10mM Tris-|(2-carboxyethyl) phosphine hydrochloride (TCEP) (#148415000, ACROS Organics)/40 mM 2-chromoacetamide (CAA) (#C4706-2G, Sigma) for 5 minutes at 70°C. The samples were digested overnight at 37°C with 1 μg trypsin (sequencing grade; #V5111, Promega) per 100 μg of protein.

### Chemical cross-linking coupled with mass spectrometric analysis

LC-MS was performed using Ultimate^®^ 3000 HPLC series for peptide concentration and separation. Nano Series ™ Standards Columns were then utilised to separate the samples. A linear gradient from 4% to 25% solvent B (0.1% formic acid in acetonitrile) was applied over 30 min, followed by 25% to 90% solvent B for 20 min. Peptides were eluted using at a rate of 250 nL min^−1^ using a Triversa Nanomate nano spray into the Orbitrap Fusion mass spectrometer (ThermoScientific). Mass scan range of 375-1500 were used for the peptide precursors at 120 K resolution, with automatic gain control of 4×10^5^. Precursor ions range of 2-7 were isolated and fragmented using Higher-energy Collisional Dissociation (HCD) fragmentation using the Orbitrap detector at a resolution of 30 K. MS/MS fragmentation was performed using a collision energy of 33%, with a maximum injection time of 200 ms and automatic gain control of 1×10^4^. Dynamic exclusion duration of 45 s with 5 ppm tolerance was used for the selected precursor and its isotopes. The instrument was run with a cycle time of 2 s. 20 ul of the samples were injected into the nano LC-ESI-MS/MS using an Ultimate 3000/Orbitrap Fusion (Thermo Scientific) using a 60-minute LC separation over a 50 cm column. The ProteoWizard MSConvert toolkit^42^ was used to convert the raw data files into .mgf format. Scaffold Proteome Software was used for sequence visualization and coverage. Cross-linked peptides were analysed using the Stavrox software^110^, using the in-built parameters for either BS3 or EDC. Precursor and fragment ion tolerance were set to 10 ppm.

The spectra were manually inspected, and continuous fragment ions were expected to be seen for both peptides. Cross-linked peptides were identified in two replicate datasets. Detected peptides were listed in Table S4.

## ACKNOWLEDGEMENTS

We thank Dr. Masayuki Onishi (Duke University) for critical reading and constructive comments on a early version of the manuscript. Computational analysis was facilitated by resources provided by the Swedish National Infrastructure for Computing (SNIC) at the Uppsala Multidisciplinary Center for Advanced Computational Science (UPPMAX), partially funded by the Swedish Research Council through grant agreement no. 2018-05973. We thank the Warwick Proteomics RTP for mass spectrometry. MKB was supported by the Wellcome Trust (WT101885MA) and the European Research Council (ERC-2014-ADG No. 671083). Work by the NR laboratory was supported by start-up funds from the Division of Biomedical and Life Sciences (BLS, Lancaster University) and a Leverhulme Research Project Grant (RPG-2019-297). NR would like to thank Johanna Syrjanen for performing trial expressions of the Odinarchaeota ESCRT proteins, and Joseph Maman for helpful discussion regarding the SEC-MALS. NR, WX and AP would like to thank Charley Lai and Siu-Kei Yau for assistance with initial Odinarchaeota ESCRT protein purifications. DPS and BB would like to thank Chris Johnson at the MRC LMB Biophysics facility for performing the SEC-MALS assay on Heimdall Vps22. TH, HH, MB, RS, JL, D Tamarit, TE, DPS and BB received support from a Wellcome Trust collaborative award (203276/Z/16/Z). BB and DPS were supported by the MRC. DT was supported by the Swedish Research Council (International Postdoc grant 2018-06609).

## AUTHOR CONTRIBUTIONS

TH, BB, NR, MB conceived and integrated the overall study.

TH, SP, NR, DPS, D Tamarit, RS made all figures.

TH, MB, NR, DPS, D Tamarit, and BB wrote the first draft of the manuscript All authors contributed to reading and revising the manuscript.

TH (under supervision of MB): carried out all Heimdall experiments in Figures 2 and 3 (except for SEC-MALS: DPS- and the analysis of mass spectrometry-HH), all iTASSER modelling of Heimdall proteins, and helped make Y2H plasmids, and carried out preliminary phylogenetic analysis.

SP (under supervision of MB): made most plasmids for Y2H and carried out the entire pairwise Y2H screen.

DP (under supervision of NR): Expressed and purified the Odinarchaeota ESCRT proteins (with AH, WX, AP, MM and NR), performed the CD analyses (with MM and DT), the Odin SEC-MALS analyses (with NR and DR) performed the ESCRT-II molecular modelling (with NR) and designed and purified Vps25 constructs for crystallography (with NR).

DPS (under supervision of BB): Expressed, purified, and carried out the analytical gel filtration on Heimdall Vps22, collected sequences for phylogenetic studies and helped in the bioinformatics analyses of D Tamarit.

RS (under supervision of JL): solved the structure of Vps25

D Tamarit (under supervision of TE): performed all genomic and phylogenetic analyses.

HMAH: Processed and analyzed XL-MS data.

**SP, DP, DPS, RS and D Tamarit contributed equally to this work and are listed alphabetically.**

