## Supplemental Tables 1-4 for "Characterisation of the Ubiquitin-ESCRT pathway in Asgard archaea sheds new light on origins of membrane trafficking in eukaryotes"

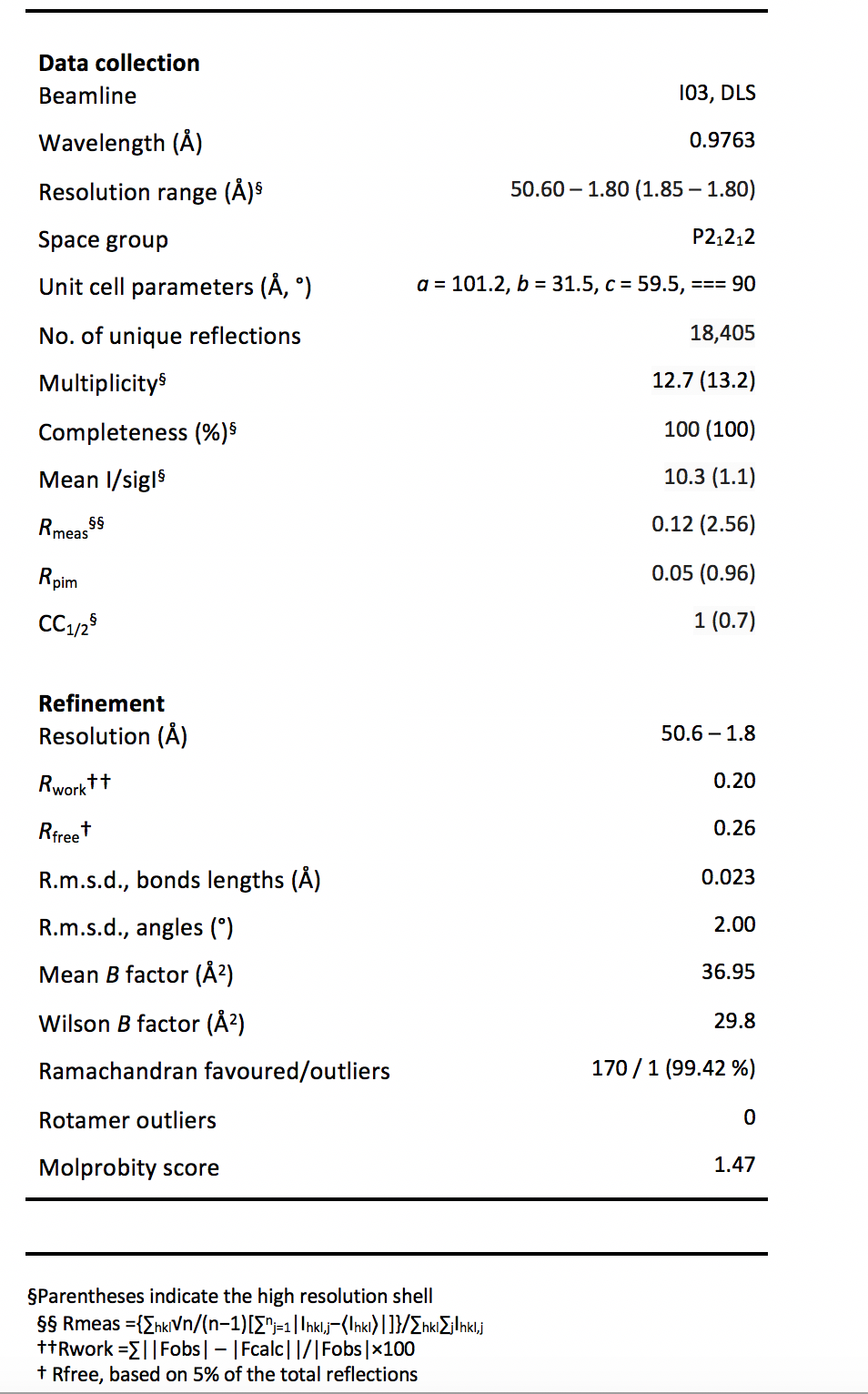

Supplementary Table 1. Data collection and refinement statistics for the Odinarchaeota Vps25ΔN crystal structure.

Table 2: Predicted amino acid sequences of Asgard ESCRT proteins used in this study

>Lokiarch_37670|Ub

MTQYRFRLTGVPPDQAIKLIELDPNNAIADIKKTVQREYKLNPILAIQFIFKGKVLPDQLKFAKIGIHPKKDVITVMATQAGGISIIH

>Lokiarch_16100|Vps23

MDKQRILREAQKIAKKFSFWMVSGNIAHLYGYVYEIEEKKFELEIKFDENFPTNPPQFIYQKDIIELLENVQLDKLINWTQESDVVDIIDELKMKIQKVLHVPKTILEENIKLKEVSENSNGSFEIDNFITPDLSRYPPDSQYNEYLTPSDPNNNSFYDEQSVKPSPGEPQQSFENSSNTPTKPIQENFFEESSQMNVTLNTELSLIQREYTYDQISEKSAEINIYITITLTKTFIIGVDFTNYPKKPIFHFPQEVKKLLGDPQKSLNILRTWNPKKPLHIVDILHELEKNLYFFKEIELQFKEITGEYQYEAVSESLTALKVSILTYGFKDYKLNVDLKTYPNPPIINFTSELQQLINIPTTELASFKNWINKKSKVVEIIREISWLVDKNSRINFEIDLLKDHYKDMKYDPIYSTLCVNMKGKMKTEDLTFEFQIILPVEYPMKIPEIKVLNEFELETHEKIKKDLYNSFDDFFNDWTPFSYLIDLFNLISK

>Lokiarch_10170|Vps28

MADFNTNTIKDENIESKLFALIHTILNAFQKFEEGLINDTFFRKTVKNAIKSLIKINFYLNEKRIKLSHLLNKMNFTSHYNDAITIINTITSKEISNISIENNFSESTQSSKEELSSILMELPGITLEITSSFITLMDALKLKGINEGELIIKMFKTLIKNVKRFPGNEEIHFNLKQIYKHVLSYMDREEGIDGFNEKIVDELYQIFKDFQRKLNLKR

>Lokiarch_37450|Vps22

MGIRKIEDEIDLMEAMKKKAAELKIKDLEARASVERFTKDQLRELAKKHGILLALTPELRKEVKKLQKKYNIPSSKIITEQIIEHDVLKKFSKIDHRKLGMLAYQRVLMSKEETGGIIPLSEVYELVNTGILKSNIEVKDVEKAMRLLRKMRIIEGLTELSSGSMLVRFFPIQFTGDEAKVIELAKEKGLLTLENVCVMLKWSQDRALRALESLEKSGVAKFRENILTGKKWFFPSL

>Lokiarch_37460|Vps25

MERQRERQEELPKDVKEYTEKGLDELKKELEKLKNLKSEKEQLEKEAAEKNIIKEEDFKIADLSYIEEQFDKLERVLNSQIGDIDSKTYKRYAEQIESGLQELEEEIIGEKGLIEKKITAHEKLINAYPWLEDERKRFMYTLPDKNKQKADYTSWKIEWAKVLFDYARFAILHTIYIRELNSQKPFTDFTNREQYILEIVDELIAQKQAKWFSKKKEKLRVYWKTLEGWSDEIYKWAYENGKLEPIMIFELREAKQEDFSSLPIEDLEAIFKILAKSRRAKILKLENGQLAFKIKLE

>Lokiarch_16760|ESCRT3A

MGLKKKLFPRKKGKEKANLITNAKVHIHKLNLVNRNYTKRAEISRKNAKIALRRGEKTRAKNFLIQYKSYNAKIDRSNNIRSKIERQIQAIEEGQLISQTGSIFEGIRDELKYIATEASPAKVAEIAEDSDVYVSEIEEAADILAGDPEIDLGIDVTDELNQLETELLLSQGGTMPDAPSDDLQYIPEYGDELEEEVESKTKEKVQAEIEKLRKELES

>Lokiarch_37480|ESCRT3B

MGFLKDISKWLSGGGKKIDSVRSASIKLKVFNKRLMRQTKKLEITAKIARDKAVNLRKQGDLDGSKFHARNYLQVKKQARAVEFFRTNLEGLQFKLDQANAVKDVSQIMTGIAQSLSVLKNQLSVPQIQEVMSQIDIDMEEFAITADITTEGMDSISIDTAVTDSDVNEVLGEMDAEIQVEMGAELPTAASDEKIAELEKELNRLKSDE

>Lokiarch_37470|Vps4

MSSVDSKLYQFAVSKAKEAVALDSQGKHRQAINSYLRAAEILVQFMKFNKNPQMRSLCQRNIDDYLARAKILKSQLGGSGSRRSRPTGKSGISKGGDSSTTTNGEVSSEEQELIDMISGTIVTESPNVRWGDIAGLENVKQALREAIVLPIIKPELFKGARKPWSGILLFGPPGCGKTLLARAAATECKATFFSASSADLLSKWLGESEKLISSLFKVARLKAPSLIFMDEIDSIATKRGEGSESGGERRVKTQLLSEMQGLRSTYDKPLLVLGATNRPWDIDNAMLSRFEKRVRVPLPDLKARSGICKIHTEGINTSLTNEDYVELGVRTEGYSGRDIANLCREVIMLPIRELDTKGLLENSDQEITLRDIDIKDFTKTLKKVKPMTSKSLMKQYNEWASEFGE

>ThorAB25_08990|SB

MCEDCGAALCSECLESRSSEYTVCSDCHHTLGSPMPGEKFEECPQCESKELSKGRKTVDICPRCHSSRVVFLEDRRRTLALDMRHAIMSIQYGHTKLREFANKLAGSKRLLVSLRMANFLHYRWIEEKIESMQSELPAIKNRISSQAEIIAKRMAAETKDFIDYTKWNPTQFPFIEGITNRITEMGIYYKQNVDESLESLRGTLKDVRRQLDGLDYYKGQFTAFYEYGQLSVGELPVCALPEIKITGSDFLKNDKAFGTLFITNKRLLFIAETGRVRKKMETVFDFPLIYLKSIEEDGRLRKRVVLKLKQGEIKASCSDQTKKVLPDYIEIARKFEKYIQTDMQRVRKIEQADISTSDVRLKIEGMVYSLLSTNDRSQYNEEMPLMRPVPRPRVGDWKSTPIRHYPYDGYYRKPESFHDELDRVIGQYDQRPSARPTYDASPRVHSLQRDAASIENAIRDAVHMLRSGQLAAEDFIRRYRDLMRDSYHNRREIERVTRETRDKLW

>ThorAB25_09030|Vps22

MHERLHSELFASIATLDQLKKLYDLGAVDNRSYQRESQALLHDVRSQIENMKRIGLDIGTFVASERIIEAYPEAATLLRLGGLAPRRSLQGMTLDQFYDYIGLAILDVATEHSSKGLMGLAELIVQVVRRLPEIESVTYNDVIEAVNRVAENGLIPGIRTLRGGVKVVELRPLELRRDQADVINLAASKGYVSVEEVIMQLKWSEDHARQVLSSLVEAGMAVADIRFSTGGRYYFPGLRARDNSDPRHTGAY

>ThorAB25_09020|Vps25

MGVRERKKEKTKTKESTEPAEQEQVESSPDVKETITEEEVHTPEVVVEVAEAVETMEPEKQAEAVNETVSEVVVAEPDFVMPSWGDVEETEWMYGIPTREEDRTLWAEEWSDYLLEWTKE

KKVHVLSLATIISEPPFKDLRNKVDSFKAIAAVLIDKEVAEWVDKKKRQLRVYWRPLEDWVDILYQWALKTGKLRLDVKSIVIQEIDEDFATLPEKDIQIVLALMVKKKLAEWIDEKRGAILIST

>ThorAB25_09000|ESCRT3A

MRFLGSIRKAFSWGRKKPDIGEISSRLRILSKQLARQRNKLEKEERDTKARAVRARKAGQTEAYRTYATEMVRFRRYALSVDKSRLQILKILAHLNRAQTSAKTQKALQEVAKILGMLGDGSDASKVVANVDEIARRLEEFEIEASISDEALDSTMEITSEDLSSAFQEIDAEAGLADAPIATRPAGEADSLEEDIKALERELGV

>ThorAB25_07240|ESCRT3B

MGFLSGKKKVDAERGILEMKGTMNALTLRVRRLETRMTEEQKSAKDSMSQGNRSQARSHLSIVVDLENRRSRYQQQFLTLETALLNLEEAKTQAEVLRAFTIANDALTEARSMLKPEEIQTQLDKLSESFEHISIAGELLSEDLSGADSTIADEERIDRQLESMEAEVLLEKEGVLPPLEAEGAITTKTSKTETSDEKRIDELLADLEKEARDERAREKES

>ThorAB25_09010|Vps4

MADRVDHGLKTLLAKAVELDQRGMKKDACQEYLKASKILIKLSNSAALPSVKKHYLDRAQECVDRVRYLSGIKKMAHGPVDGLVAPPKVDVGTTEPVPIQETSKAEVELDDEEKRLREMISDTIITERPNMKMSEVAGLADAKQAIDDAIVVPMKHPELFKGKARQPWRGILFYGPAGCGKTLVAKAVASEVNATFFNVSAANIVSKWLGESERLVMNLFGLARKNQPAIIFIDELDSIGVSRSGDDVGGERRLKTQLLTELQGLASNEEDRITLIGATNLPWELDFALRSRFEKKIHVPLPGKEARAKIFEIHMEDVEVSPTVEYEELADLTEGYSGRDISVVCREAAMEPIRDLQRTGRMDDEQEILDIRPVSRDDFLHAIENIRPATPPEDVKRYIDWAEGS

>HeimAB125_14240|Ubiquitin

MRITVVTAIGGGKIDLDVDPHSTVAQLKREVAKKKKIPANTVIIVFHGKQLDDSEILKQTGMMDGDKCYLIARTKGG

>HeimAB125_14070|Vps23_Vps28

MNKISTDDFLLANEMQLVFDKQLNFEPVRDNLKQWKGYVGTIEDTEIPIFANVYIQDGFPDVPPFVDITPKVNHPNVDEDGTLSMRLLAEWEPRYHIYQVMIDIKQLFINVPAKTYKTKIKKPKEEQRYTGLPRATIPVRTLTQVSLTGIDPEIKEEKLSIDSSIMYYQNQIEQVTRDIEKERESLLKEGGIAVQSRTKEISISREEDLRAESIAVQDTLEVIQEKFEDGDMTNIDFLKLHKRYTKKGFKAEKQLNIIKSSSSGDMTKVEEKILNLEADLFASIATLDNLVRSYENSDIEQIAYRKQLRSLIRGIFKTRLELEKVGFNLDDFVKRENLNNRYPKGIKQLMVVEGEETADTNSIPFDTLKKMPSKTADFVSSAIELIDLTRLRSVARADLLLADIEEMIHILTTFPSIPKDHWVIGDMNNWRDIIGKYKTQEIIKEEDCEKLEFQAARWLNDFRRVLKEL

>HeimAB125_14050|Vps22

MGLRDIEKKIKVKSAFQVKEAELMIKELEQMIDNISEVKKQLKKFEKKHGDKISENKDYYEKLSNLREELGLPTEIGRFDWKEAPTLKDKLTGKGFYDILANELLEIGQQLIPDNGGIMAIGELFTQLNKIRPGRMVSIDDMYRSLDKLINSQLIPPYRVLDSGLKIIEFVPVEFSPDHDVILNYASRVGFVTKEDLLMRTSWTEERIDRCLKFFEDNSIARVDRSYSEGVKFWFPGLMGL

>HeimAB125_14040|Vps25

MSTTEKKKPKKEKSSEIDLKKLKKEKEISDEKLKLIEASIKGLEITAEELKSTLPVDAIIIKEDLSKDQRAVLELEFEDELPAWVKRPWMYINPNKADQVESWLQSWSAIVLDYSRIFVIHIININDLRSDHPFNNKKLGKRLTLAQVRSVIDHLVQQNVARWLDEDTKTRARIYWKSNEDWADALLDFMIETGRVVEVHSLYDLTQMDQDWGVLPADDLIIVCEMLVERKVARWLNKEKTIIQFEADKVF

>HeimAB125_14010|ESCRT3A

MGWLLGTKKSPRDHIINLRLTVRRLERAQRKLARDESKMQLKMKQSIQRGDLESARLFATDIVRSRRWEFGYQKLVSRMNGLIFKLERADTAASMAEEMHGIASALRAANAQLQIPDLDRVIQDMESSIDGIEESTGTIEDGIDDLLVADSDPAEVDRLIEQTAAELGVSTQAGLPTVGVVQTDDLEEEIQKLRRKEED

>HeimAB125_14020|ESCRT3B

MVKNWLFGKKRKEDADALATLKGQQNRLQAEARNLERQSDEQKILASKMLKAGNKAGARQALKRRAVFMKRLNTVHNTAMNLQAQIDSIQTATSTAETVKAMELGTKVVGEKIKTVSPERTERVMDSVMEQRDQIEMMTEALSDPSLSEGILDFEDDAAIDEQLAQLEAEMDLGTTTSLPDVSGLPSTPVGTGEKEEDTSELEAELEGLKKKMSEDKQ

>HeimAB125_14030|Vps4

MSAQGLIDHARDFAVKAYSFDKEKNYSEAIPHYLDAAEALMKAIKFERNPQVAGSLKKKANMYISRAKELKELLKKRKERKSTAGNDEEEEKLESAISDIIVTEKPNITLNEIAGLENAKQTLREAIVLPLMRPDLFSGARRPWKGILLFGPPGCGKTLLAKATAAEVEATFFNVSASSIISKWLGESERLVKQLFELADEKQPSIIFVDEVDALAGARGGEHDAMRRVKTELLTSMDGLSSSETDRIVTVGATNMPESIDAAFRRRFERRIYIPLPDLPARGAIFLLNSKGVDLAEDVDFEVLAEITEGYSGSDIAMVCREAIMTPIREMDMAGAIGDTSIMARSATQDDYLEAIESINPSVSDDEVEKYDNWNEEFGSSA

>OdinLCB4_14240|Ubiquitin

GSHMKIEIVSAIGGEKTILDVKPETTFKEIKEIMSQKRNLQADNFVLAFRGREQEDHITLKEAGVSEEDRVYLITRTEGGSVKY

>OdinLCB4_14270|Vps23(TSG101) [ESCRT-I]

GSHMDTSIIYQEAELMYKMAPGFKTVNGDLRNWIGEIKFKSGRDFSSCWIEILLPENYPSEPPIVSCSSSLRHPIIDPATNRFNLRLLRNWNPNTHIYEIVNVIKSEFSRQPPEFLGALKPITKPSLEVENLSAGEIKELESKIRELESEVEYLRKDLAEKNDEIARLQNILSKETSFKDKGVKSGLSSLPVDSSTDLEGEKAALEGVIRDLEARFEAGDITPEEYFKLSQKYRKQLFLVGKNLSDTRKAESKN

>OdinLCB4_14280|Vps28 [ESCRT-I]

GSHMQEKNCVKESEERSRLLREEELLSSVDKETKIRIYELESELYANLATLQSISKSFEESRLDPVTFKRLLKALMKTCFKVKQELEKYGLDVKEFIKNQGFLEEFNLAIKNLEWDSNLDSSIYAISLESPGRIAVKTYELASSFITLSDSVKLRISCEVLYNLLNELFNELTKYPGFSSTHPVCKEVREWQVKLKDFNPSDILDEKTSLELENAIQTWRNEFECMIKKACM

>OdinLCB4_14290|Vps22(EAP30) [ESCRT-II]

MGISEIENRLKTKRAYTLKSIELRLIELVSNPSYKKNPELILERLFEEFGVEIFEDSSLIENLSAALKNLNIEGERLFKGLIPDISVFDKSFLGKLKVNHKKLGELLFRRASLLKRVRGGLFTVGELLDWFNRAAKYKIKVEDILKAVRILEKNRIIPGRRIIGGNVLVIQFLPLELSSDHLFILNLASSKGWVTVEEASMNLNWPLERVTLVLDKLVDYGVARLDSSYAHGKKYYFPAFLSKLEHHHHHH

>OdinLCB4_14300|Vps25 [ESCRT-II]

MELTKRRDKNTLKKEELLEQINLLKKELDAEKTQVEGVGVAGRVKQEEMITGKADTQILTVGFEWAAQPEKYPWMYSLPSKTEDFEDWLNQWSDFTLQWFKINKLHQISLVELMGEKPFSYLQNKSKALTVIVENLIARNFCKYTDKEYKSIRVFWRGYRDWSEVIYNWALKKGRTELTFFEIIDLKESPDNFHMLPKEDFKKIFNILVKNKRAEWINKKNMHIRILFLEHHHHHH

>OdinLCB4_14300 Truncated|Vps25ΔN [ESCRT-II]

MLTVGFEWAAQPEKYPWMYSLPSKTEDFEDWLNQWSDFTLQWFKINKLHQISLVELMGEKPFSYLQNKSKALTVIVENLIARNFCKYTDKEYKSIRVFWRGYRDWSEVIYNWALKKGRTELTFFEIIDLKESPDNFHMLPKEDFKKIFNILVKNKRAEWINKKNMHIRILFLEHHHHHH

Supplementary Table 3:

| ***OdinLCB4 Y2H plasmids*** | | |
| --- | --- | --- |
| Collection # | Vector | Gene |
| pSPW378 | pMM5 | **Ubiquitin** |
| pSPW366 | pMM5 | **Vps23** |
| pSPW334 | pMM5 | **Vps28** |
| pSPW344 | pMM5 | **Vps22** |
| pSPW380 | pMM5 | **Vps25** |
| pSPW354 | pMM5 | **ESCRT-III (A)** |
| pSPW360 | pMM5 | **ESCRT-III (B)** |
| pSPW368 | pMM5 | **Vps4** |
| pSPW430 | pMM6 | **Ubiquitin** |
| pSPW418 | pMM6 | **Vps23** |
| pSPW386 | pMM6 | **Vps28** |
| pSPW396 | pMM6 | **Vps22** |
| pSPW432 | pMM6 | **Vps25** |
| pSPW406 | pMM6 | **ESCRT-III (A)** |
| pSPW412 | pMM6 | **ESCRT-III (B)** |
| pSPW420 | pMM6 | **Vps4** |
| ***Lokiarch Y2H plasmids*** | | |
| Collection # | Vector | Gene |
| pSPW376 | pMM5 | **Ubiquitin** |
| pSPW364 | pMM5 | **Vps23** |
| pSPW332 | pMM5 | **Vps28** |
| pSPW336 | pMM5 | **SB** |
| pSPW342 | pMM5 | **Vps22** |
| pSPW348 | pMM5 | **Vps25** |
| pSPW352 | pMM5 | **ESCRT-III (B)** |
| pSPW358 | pMM5 | **ESCRT-III (A)** |
| pSPW372 | pMM5 | **Vps4** |
| pSPW428 | pMM6 | **Ubiquitin** |
| pSPW416 | pMM6 | **Vps23** |
| pSPW384 | pMM6 | **Vps28** |
| pSPW388 | pMM6 | **SB** |
| pSPW394 | pMM6 | **Vps22** |
| pSPW400 | pMM6 | **Vps25** |
| pSPW404 | pMM6 | **ESCRT-III (B)** |
| pSPW410 | pMM6 | **ESCRT-III (A)** |
| pSPW424 | pMM6 | **Vps4** |
| ***HeimAB125 Y2H plasmids*** | | |
| Collection # | Vector | Gene |
| pSPW374 | pMM5 | **Ubiquitin** |
| pSPW362 | pMM5 | **Vps23** |
| pSPW330 | pMM5 | **Vps28** |
| pSPW340 | pMM5 | **Vps22** |
| pSPW346 | pMM5 | **Vps25** |
| pSPW350 | pMM5 | **ESCRT-III (A)** |
| pSPW356 | pMM5 | **ESCRT-III (B)** |
| pSPW478 | pMM5 | **ESCRT-III (B) 52C** |
| pSPW370 | pMM5 | **Vps4** |
| pSPW426 | pMM6 | **Ubiquitin** |
| pSPW414 | pMM6 | **Vps23** |
| pSPW382 | pMM6 | **Vps28** |
| pSPW392 | pMM6 | **Vps22** |
| pSPW398 | pMM6 | **Vps25** |
| pSPW402 | pMM6 | **ESCRT-III (A)** |
| pSPW408 | pMM6 | **ESCRT-III (B)** |
| pSPW478 | pMM6 | **ESCRT-III (B) 52C** |
| pSPW422 | pMM6 | **Vps4** |
| S.pombe_ESCRT-Y2H | | |
| ***S.pombe Y2H plasmids*** | | |
| Collection # | Vector | Gene |
| pSPW738 | pMM5 | **SpUbi4** |
| pSPW739 | pMM5 | **SpSst6** |
| pSPW740 | pMM5 | **SpVps28** |
| pSPW741 | pMM5 | **SpVps36** |
| pSPW742 | pMM5 | **SpVps22(Dot2)** |
| pSPW743 | pMM5 | **SpVps25** |
| pSPW744 | pMM5 | **SpDid2** |
| pSPW745 | pMM5 | **SpVps20** |
| pSPW746 | pMM5 | **SpVps32** |
| pSPW747 | pMM5 | **SpVps1(Did4)** |
| pSPW748 | pMM5 | **SpVps24** |
| pSPW749 | pMM5 | **SpVps4** |
| pSPW750 | pMM5 | **SpCmp7** |
| pSPW751 | pMM6 | **SpUbi4** |
| pSPW752 | pMM6 | **SpSst6** |
| pSPW753 | pMM6 | **SpVps28** |
| pSPW754 | pMM6 | **SpVps36** |
| pSPW755 | pMM6 | **SpVps22(Dot2)** |
| pSPW756 | pMM6 | **SpVps25** |
| pSPW757 | pMM6 | **SpDid2** |
| pSPW758 | pMM6 | **SpVps20** |
| pSPW759 | pMM6 | **SpVps32** |
| pSPW760 | pMM6 | **SpVps1(Did4)** |
| pSPW761 | pMM6 | **SpVps24** |
| pSPW762 | pMM6 | **SpVps4** |
| pSPW763 | pMM6 | **SpCmp7** |

Supplemental Table 4: Mass spectrometry identifying intra vps22 crosslinks

| **Raw File** | **Peptide 1** | **Peptide Sequence 1** | **Cross-link residue 1** | **Peptide 2** | **Peptide Sequence 2** | **Cross-link residue 1** | **xlink type** | **xlinker used** | **Stavrox Score** | **M/Z score** | **charge state** | **Additional information** |
| --- | --- | --- | --- | --- | --- | --- | --- | --- | --- | --- | --- | --- |
| MS19-237_VPS22_BS3_DCL | DKLTGK | 89_94 | 90 | SLDKLINSQLIPPYR | 145_159 | 145 | intra-molecular | BS3 | 241 | 639.617 | 4 | (+3 form found in MS19-241 BS3 DCL1) |
| MS19-237_VPS22_BS3_DCL | MGLR | 0_4 | { | IKVKSAFQVK | 10_19 | 13 | intra-molecular | BS3 | 193 | 441.016 | 4 | (+3 form found in MS19-227 BS3 DCL) |
| MS19-237_VPS22_BS3_DCL | DKLTGK | 89_94 | 90 | LSNLREELGLPTEIGRFDWKEAPTLK | 63_88 | 86 | intra-molecular | BS3 | 145 | 953.523 | 4 | (same form found in MS19-227 BS3 DCL) |
| MS19-237_VPS22_BS3_DCL | DIEKK | 5_9 | 8 | VKSAFQVK | 12_19 | 13 | intra-molecular | BS3 | 137 | 639.719 | 3 | (+2 form found in MS19-227 BS3 DCL) |
| MS19-237_VPS22_BS3_DCL | KQLKK | 40_44 | 40 | IKVKSAFQVK | 10_19 | 13 | intra-molecular | BS3 | 134 | 643.744 | 3 | (Same form found in MS19-227 BS3 DCL) |
| MS19-237_VPS22_BS3_DCL | QLKK | 41_44 | 44 | KFEK | 44_47 | 44 | inter-molecular | BS3 | 176 | 666.917 | 2 | (same form found in MS19-227 BS3 DCL) |
| MS19-241_VPS22_BS3_DCL | LSNLR | 63_67 | 64 | VGFVTK | 189_194 | 194 | intra-molecular | BS3 | 108 | 695.409 | 2 | (same form found in MS19-227 BS3 DCL) |
| MS19-237_VPS22_EDC_DCL_1 | DYYEK | 58_62 | 48 | IKVKSAFQVK | 10_19 | 13 | intra-molecular | EDC | 360 | 616.008 | 3 | (same form found in MS19-237 EDC DCL2) |
| MS19-237_VPS22_EDC_DCL_1 | DKLTGK | 89_94 | 89 | SLDKLINSQLIPPYR | 145_159 | 148 | intra-molecular | EDC | 348 | 600.597 | 4 | (+3 form found in MS19-237 EDC DCL2) |
| MS19-237_VPS22_EDC_DCL_1 | IKVKSAFQVK | 10_19 | 13 | ISENKDYYEK | 53_62 | 58 | intra-molecular | EDC | 320 | 806.442 | 3 | same form found in MS19-237 EDC DCL2 |
| MS19-237_VPS22_EDC_DCL_1 | HGDK | 49_52 | 51 | IKVKSAFQVK | 10_19 | 13 | intra-molecular | EDC | 307 | 528.979 | 3 | Same form found in MS19-237 EDC DCL2 |
| MS19-237_VPS22_EDC_DCL_1 | QLKK | 41_44 | 43 | EELGLPTEIGR | 68_78 | 68 | intra-molecular | EDC | 286 | 570.996 | 3 | Same form found in MS19-237 EDC DCL2 |
| MS19-237_VPS22_EDC_DCL_1 | EELGLPTEIGR | 68_78 | 68 | SLDKLINSQLIPPYR | 145_159 | 148 | intra-molecular | EDC | 281 | 738.661 | 3 | Same form found in MS19-237 EDC DCL2 |
| MS19-237_VPS22_EDC_DCL_1 | KFEK | 44_47 | 46 | KQLKK | 40_44 | 43 | inter-molecular | EDC | 251 | 392.92 | 3 | same form found in MS19-237 EDC DCL2 |
| MS19-237_VPS22_EDC_DCL_1 | DKLTGK | 89_94 | 89 | KFEKK | 44_48 | 47 | intra-molecular | EDC | 194 | 661.396 | 2 | same form found in MS19-237 EDC DCL2 |
| MS19-237_VPS22_EDC_DCL_1 | VLDSGLK | 160_166 | 162 | VLDSGLK | 160_166 | 166 | inter-molecular | EDC | 119 | 722.425 | 2 | same form found in MS19-237 EDC DCL2 |
| MS19-237_VPS22_EDC_DCL_1 | DIEKK | 5_9 | 7 | VKSAFQVK | 12_19 | 13 | intra-molecular | EDC | 139 | 507.3 | 3 | similar form found in MS19-227 EDC DCL |
| MS19-237_VPS22_EDC_DCL_1 | DKLTGK | 89_94 | 90 | LSNLREELGLPTEIGR | 63_78 | 68 | intra-molecular | EDC | 205 | 813.79 | 3 | same form found in MS19-237 EDC DCL2 |
| MS19-237_VPS22_EDC_DCL_1 | VKSAFQVK | 12_19 | 13 | ISENKDYYEK | 53_62 | 55 | intra-molecular | EDC | 179 | 726.049 | 3 | same form found in MS19-237 EDC DCL2 |
