## Supplemental Figure Legends for "Characterisation of the Ubiquitin-ESCRT pathway in Asgard archaea sheds new light on origins of membrane trafficking in eukaryotes"

**Figure S1: Schematic representation of the Eukaryotic ESCRT pathway.**

Ubiquitin is recognised by the UEV domains of Vps23 (and some Vps37 homologues) of the Vps37/Vps23/Vps28 ESCRT-I ternary complex. The C-terminal alpha-helical bundle of Vps28 interacts with the GLUE/zinc finger domain of ESCRT-II to bridge the ESCRT-I and -II subcomplexes. The eukaryotic ESCRT-II complex in yeast and human systems forms a Y-shaped tetramer consisting of a Vps22/Vps36 stalk leading to a splayed Vps25 dimer. The core of all the related ESCRT-II subunits consist of tandem WH domains. The splayed Vps25 dimer then recruits the ESCRT-III membrane scission machinery with a geometry that facilitates the formation of spiral ESCRT-III filaments. The pathway start at the bottom of this schematic, with ESCRT I / Ubiquitin interaction and ends at the top of the schematic, with ESCRT III mediated membrane remodelling.

**Figure S2.** **Phylogenetic classification of Asgard archaeal bins into robust lineages using 15 co-locating ribosomal proteins.** Genomes downloaded from public databases were reannotated and searched for ribosomal proteins from a set of 15 encoded in the same gene cluster. Sequences of 5 or more of these ribosomal proteins encoded in the same contig were aligned, trimmed, and concatenated for phylogenetic reconstruction. A maximum-likelihood phylogeny was reconstructed using IQTree under the LG+C60+R4+F model. Support values represent approximate likelihood ratio tests (left) and ultrafast bootstrap (right) based on 1000 pseudoreplicates each. Outgroup sequences were all non-Korarchaeal Crenarchaeota species representatives available at GTDB on December 5th. Only genomes confidently classified as Heimdall-, Hel-, Loki-, Odin- and Thorarchaea (colored) were used for further analyses.

**Figure S3. Synteny plot of ESCRT-Ub genes in Asgard archaea.** Maps around genes containing ESCRT and Ubiquitin domains for Heimdall- (A), Thor- (B), Hel- (C) and Lokiarchaeota (D), using GenoPlotR. Arrows represent genes and are colored if their products were annotated as containing diagnostic domains for Ub/ESCRT proteins (see legend in A). Genome regions are plotted at 10 kb of ubiquitin or ESCRT protein-encoding genes (colored), or until a contig boundary (thicker vertical lines). Similarity lines indicate best-reciprocal BLAST-p hits with an e-value lower than 1e-5.

**Figure S4. Phylogenetic reconstruction of ESCRT-II genes.** (A) Unrooted maximum likelihood phylogenetic tree of Vps22 (blue), Vps36 (purple), Vps25 (orange) and outgroup (black) sequences corresponding to the same dataset as Fig 1D. The tree was reconstructed using IQ-Tree under the LG+C60+R4+F+PMSF model. Support values represent transfer-bootstrap expectation (left) values or standard Felsenstein bootstrap proportions (right) based on 100 bootstrap pseudorreplicates. (B) Unrooted maximum likelihood phylogenetic tree of the same sequence dataset as (A) without outgroup sequences. The tree was reconstructed using IQ-Tree under the LG+C60+R4+F model. Support values represent approximate likelihood ratio tests (left) and ultrafast bootstrap (right) based on 1000 pseudoreplicates each. (C) Unrooted maximum likelihood phylogenetic tree of the same sequence dataset as (B) plus three small outgroup clades included in (A) (black), and all homologs identified through a Psiblast search against NR (see Methods) and classified as neither eukaryotic or Asgard archaeal (black). The tree was reconstructed using IQ-Tree under the LG+C20+R4+F model. Support values represent approximate likelihood ratio tests (left) and ultrafast bootstrap (right) based on 1000 pseudoreplicates each.

**Figure S5. ESCRT/Ub domain detection.** Number of genes with protein domains related to ESCRT and Ubiquitin metabolism in Asgard archaea. (A) Number of genes containing specific Interpro domains and domain combinations. Parentheses include the gene families that the corresponding domains represent. (B) Number of genes containing their corresponding Interpro domains or domain combinations shown in (A). Asterisks behind “Ubiquitin” and “E2” indicate genes with domains that are characteristic of these proteins, but that, in combination with E1 and ESCRT-I domains (respectively), are likely to simply represent the latter.

**Figure S6. Alignment of E2- and UEV-domain-containing proteins.** Alignment between genes identified as Ubiquitin-conjugating enzyme E2 (black), UEV-containing Vps23/Vps37 (orange) or as a gene fusion between Vps23/Vps37 and Vps28 (red), performed with Mafft E-INS-i (v7.450) and visualized with <http://was.bi/>.

**Figure S7. A. Ubiquitin-binding enzyme (E2)-like protein found in Heimdallarchaeota.** The model structure of the E2-like domain of Heimdallarchaeota (light blue) is superimposed on the 3D-structure of budding yeast E2 domain (PDB:1JBB, pink). The catalytic Cys residue (red) in the 3D structure of the E2 domain (PDB:1JBB) and its equivalent amino acid residues (blue) in the model structure at the position are highlighted. (i) HeimAB125_07740 (27th-135th amino acid residues) ;(ii) HeimAB125_09840 (1st-107th amino acid residues); (iii) HeimAB125_14070 (1st-130th amino acid residues); (iv) HeimAB125_11700 (25th-122nd amino acid residues). Schematic domain architecture of Asgard E2L proteins were also shown. **B. Multiple sequence alignment of E2L proteins in Asagard archaea.** Multiple alignment of E2L proteins were performed by MUSCL. Alignmet for Asgard E2L were shown.

**Figure S8. Domain organisation of selected Asgard ESCRT-I proteins.** The domains found in Heimdallarchaeota Vps23/28 and Odinarchaeota Vps23 and Vps28 are compared to the ones present in *S. cerevisiae* ESCRT-I subunits. The following domains are shown (homologous regions are represented with similar colours): UEV, Ubiquitin E2 variant domain; 23-Stalk, Vps23 stalk region; 23-SB, Vps23 steadiness box; 28-SB, Vps28 steadiness box-like; 28-Nt, Vps28 N-terminal extra helices; 28-Ct, Vps28 C-terminal four-helix bundle; PRR, Vps23 proline-rich region; 37-Stalk, Vps37 stalk region; 37-SB, Vps37 steadiness box-like. The total number of residues for each protein is indicated.

**Figure S9. Size-exclusion chromatography analyses demonstrating the physical interaction between the Odinarchaeota ubiquitin homologue and the Odinarchaeota Vps23(TSG101) ESCRT-I subunit.** From Top to Bottom: Odinarchaeota ubiquitin protein only (top); Vps23(TSG101) protein only (middle); Vps23 (TSG101) pre-incubated with ubiquitin, demonstrating stable complex formation (bottom). All proteins were separated on a Superdex S200 HR 10/300 size exclusion chromatography column. The relative elution volumes of the size standards b-amylase (200 kDa), alcohol dehydrogenase (150 kDa), bovine serum albumin (BSA) (66 kDa) and carbonic anhydrase (29 kDa) and cytochrome-c (12.4 kDa) are also indicated (in grey). Eluted fractions were resolved by SDS-PAGE and visualised by Coomassie stain. Chromatography UV traces (at 280 nm) for the respective elution profiles are displayed to the left of each panel.

**Figure S10. SEC-MALS analyses of the Odinarchaeota ESCRT-I and -II subcomplex proteins.** **A.** Analysis of the Vps23(TSG101) ESCRT-I protein that has a fitted molecular weight of 60.37%(±0.91%), consistent with a stable dimer. **B.** Analysis of the Vps28 ESCRT-I protein that has a fitted molecular weight of 32.48% (±2.49%), consistent with a monomer. Note that a minor larger peak consistent with an unstable and transient dimer is also detectable. **C.** Analysis of the Vps23(TSG101)-Vps28 ESCRT-I complex that has a fitted molecular weight of 88.27% (±0.74%), consistent with a trimeric arrangement based on the Vps23(TSG101) dimer in complex with a Vps28 monomer. **D.** Analysis of the Vps22(EAP30) ESCRT-II protein that has a fitted molecular weight of 28.29% (±3.51%), consistent with a monomer. **E.** Analysis of the Vps25 full-length ESCRT-II protein that has a fitted molecular weight of 26.59% (±2.80%), consistent with a monomer. **F.** Analysis of the N-terminally truncated Vps25 ESCRT-II protein that has a fitted molecular weight of 21.59%(±1.26%), consistent with a monomer. In all panels the differential refractive index (dRI) is plotted in conjunction with molecular weight (M_w_).

**Figure S11. Chemical crosslinking coupled with MS analysis showed homodimerization of Heimdall Vps22 A.** Inter-dimeric peptide identified for HeVps22 using BS3 chemical cross-linker. **B.** Dimeric peptide identified for HeVps22 using both BS3 and EDC cross-linkers. Green colour indicates precursor ions. The red and blue colours indicate the b and y fragment ions, respectively.

**Figure S12. Structure prediction and physical biochemistry of Odinarchaeota ESCRT-II proteins.** Tandem winged-helix (WH) domains in the Odinarchaeota Vps22(EAP30) and Vps25 ESCRT-II components as demonstrated by structural modelling. (A) (left) I-TASSER (Yang *et al.,* 2014) model of the Odinarchaeota Vps22(EAP) protein (estimated TM-score 0.63) and structural superposition with a human Vps22 crystal structure (PDB: 3CUQ) (right). (B) (left) tr-Rosetta model (Yang *et al.,* 2020) of the Odinarchaeota Vps25 protein (estimated TM-score 0.39) and structural superposition with the Odinarchaeota Vps25DN crystal structure solved in this study (PDB: xxxx) (right). Also see Figure 4 for alignment of the Odinarchaeota Vps25ΔN crystal structure with the eukaryotic Vps25 homologues from *S. cerevisiae* (PDB: 1XB4) and *H. sapiens* (PDB: 2ZME).  The coiled-coil N-terminal extension unique to the Odinarchaeota Vps25 protein is highlighted in the blue circle. (C-F) Circular dichroism (CD) experimental spectra of the Odinarchaeota ESCRT-II subcomplex proteins revealing that estimated secondary structural elements, based on BeStSel fitting, agree with the predicted models (and Vps25ΔN crystal structure). Panels (C) to (E) display the experimental CD spectra results for Vps22(EAP30) ESCRT-II, Vps25 full-length ESCRT-II protein and N-terminally truncated Vps25 ESCRT-II protein (Vps25DN), respectively. In each panel the average spectra is shown for six replicate spectra (triplicate measurements for two identical samples) over a 190-250 nm wavelength range are displayed [N = 3]. (F) Summary of the percentage of secondary structural elements (% alpha helix, % beta-strand, % other) for the Odinarchaeota ESCRT-I and -II subcomplex proteins estimated from the CD spectra based on data fitting in BeStSel (All fittings had an NRMSD value < 0.1) [Micsonai et al, 2018], or from the predicted structural models (and also the Vps25ΔN crystal structure) using the STRIDE Web-server [Frishman and Argos, 1995].

**Figure S13. Y2H assay-based identification of protein-protein interactions.** **A.** The reliability of the Y2H assay used in this study was assessed using a group of known eukaryotic Ub-ESCRT pathway genes. Y2H experiments were performed using genes of the fission yeast Ub-ESCRT pathway. Positive pairs were marked with dashed line. Bait for Vps28 (*) showed autoactivation but co-presence with prey Vps25 showed stronger b-galactocidase activity. **B.** Schematic representation of interactions detected in the Y2H assays.

**Figure S14.** **Y2H assay-based identification of ASGARD ESCRT protein-protein interactions.** In this figure, interactions detected in both directions (P1-bait:P2-prey and P2-bait:P1-prey) are shown with a green filled circle and those that were observed only in one direction (P1-bait:P2-prey or P2-bait:P1-prey) are shown with a yellow filled circle.
